# Cancer drives atherosclerotic plaque vulnerability by inducing pathological angiogenesis

**DOI:** 10.64898/2026.01.18.700232

**Authors:** Lingfeng Luo, Changhao Fu, Kai-Uwe Jarr, Richard Baylis, Virginia H. Sun, Julius Heemelaar, Moritz von Scheidt, Daniela Ramirez, Johannes Krefting, Nadja Sachs, Justus Leonard Wettich, Hanna Winter, Hua Gao, Fudi Wang, Shaunak Adkar, Allen Haas, Mason Gonzalez, Kevin T. Nead, Lars Mägdefessel, Heribert Schunkert, Tomas G. Neilan, Nicholas J. Leeper

## Abstract

Cardiovascular disease (CVD) is the leading cause of death for many cancer survivors, a phenomenon traditionally attributed to shared risk factors and cardiotoxic chemotherapies. Here, we hypothesized that cancer may also directly promote atherosclerosis. Propensity-matched analyses confirmed significantly elevated cardiovascular event rates amongst cancer patients, independent of comorbidities such as smoking. Atheroprone mice implanted with colorectal tumors demonstrated accelerated features of plaque vulnerability, driven by pathological angiogenesis and intraplaque hemorrhage. Mechanistically, tumor-secreted TNF-α induced the pro-angiogenic factor LRG1 across multiple murine models and human plaques. Therapeutic interventions targeting these pathways, including with FDA-approved cytokine inhibitors or tumor resection, prevented plaque destabilization in mice and reduced coronary revascularization rates in patients. Together, these findings suggest that cancer may causally promote CVD and unveil novel translational strategies for cancer survivors.

## Introduction

Cancer and cardiovascular disease (CVD) are the world’s two leading killers, accounting for approximately 30 million lives lost each year(*1, 2*). While each disease is highly prevalent on its own, advancements in cancer therapy and cardiovascular risk reduction have paradoxically increased the likelihood of patients surviving one disease only to go on to develop the other. This phenomenon has led to a dramatic rise in the occurrence of co-prevalent disease, with nearly half of all cancer survivors expected to develop some form of CVD during follow-up(*3*). Indeed, CVD has become a leading cause of death among cancer survivors, surpassing cancer-related mortality in at least eight distinct cancer types(*4*). Furthermore, the decreasing age of first cancer diagnosis and the increasing longevity of cancer survivors have contributed to a growing burden of co-prevalent disease, especially amongst younger individuals(*5*).

The co-occurrence of cancer and CVD has historically been attributed to shared risk factors (e.g., aging, dyslipidemia, obesity, and smoking)(*6, 7*) and the widespread use of cardiotoxic therapies, such as anthracyclines and immune checkpoint inhibitors(*8*). To address these challenges, a new discipline known as ‘cardio-oncology’ has emerged which focuses on mitigating the cardiovascular side effects of cancer treatments and managing overlapping risk factors(*8, 9*).

However, emerging evidence suggests that co-prevalent disease is not solely driven by shared risk factors or treatment toxicity. Overlapping biological processes, such as clonal expansion, chronic inflammation, oxidative stress, and cellular senescence, are independently observed in both cancer and CVD, suggesting that such mechanisms may exacerbate both conditions in parallel(*10*). For example, somatic myeloid mutations in genes like TET2 and DNMT3A which predispose to hematologic malignancies are now known to have an even greater absolute impact on risk for atherosclerotic CVD(*11-13*). Additionally, unbiased genome-wide association studies (GWAS) surprisingly revealed that the top heritable locus for myocardial infarction (MI) at the chromosome 9p21 locus resides within a canonical tumor suppressor locus, further highlighting how processes traditionally assumed relevant to only one disease may have a contemporaneous impact on the other(*14, 15*).

Provocatively, recent studies suggest that one disease may also *causally* drive the other. Large epidemiological studies have demonstrated that cancer rates increase significantly following a diagnosis of CVD. This risk is particularly accentuated amongst those with atherosclerotic forms of disease, even after adjusting for traditional risk factors(*16*). Supporting this link, preclinical models have shown that inflammation following MI induces a sustained reprogramming of the immune system, and that this is sufficient to promote tumor growth and metastasis across a range of cancer models(*17-21*). Such data substantiate the concept that CVD is directly oncogenic and present opportunities for therapeutic intervention.

However, testing the causal effect of cancer on CVD has proved more challenging because the diagnosis of cancer necessitates swift, often aggressive measures to treat the cancer, and several of these cancer treatments, including radiotherapy, HER2 inhibition, anthracyclines, and T cell activators, may be cardiotoxic. Further, the pace of progress in cancer treatment means that many cardiac toxicities are only discovered after many thousands of patients receive the treatment, meaning that their cardiac risk may not emerge until after treatment paradigms have already become antiquated. Therefore, there are both known and possible unknown confounders that complicate our understanding of CVD following cancer. Here, we utilize a wide array of strategies to test the hypothesis that there is a bi-directional link underlying the rising incidence of CVD amongst cancer survivors. Defining causal mechanisms will allow for the prioritization of novel therapies to test for mitigating the burden of co-prevalent disease.

## Results

### Risk of clinical cardiovascular events following a cancer diagnosis

Prior studies suggest that cancer survivors may be at an elevated risk of developing CVD, independent of shared risk factors(*3, 22*). To confirm and extend these observations, we analyzed long-term rates of major adverse cardiovascular events (MACE) among more than one million matched individuals with and without cancer in Germany, using claims data from the statutory health insurance database within the Observational Bavarian Health Insurance Registry (Tables S1 and S2). Cancer survivors had a 25% higher rate of MACE over the subsequent nine years compared to individuals without a history of malignancy, even after adjusting for smoking history, prevalent CVD, and medication use, including statins (Fig. 1a and Fig. S1). Because this dataset could not resolve the cause of death or outpatient revascularization, we repeated an analysis of ∼400,000 individuals in the Stanford STARR database (Tables S3 and S4), who have higher-resolution data linked to the electronic health record. Consistent with the German Heart Center findings, we observed a 36% increased rate of MACE for cancer survivors (Fig. 1b and Fig. S1), including a 53% increase in acute MI (an endpoint less susceptible to confounding from anthracycline-induced congestive heart failure, Fig. 1c). Together, these data support the growing body of evidence linking cancer survivorship to increased risk for ischemic events.

**Figure 1.**
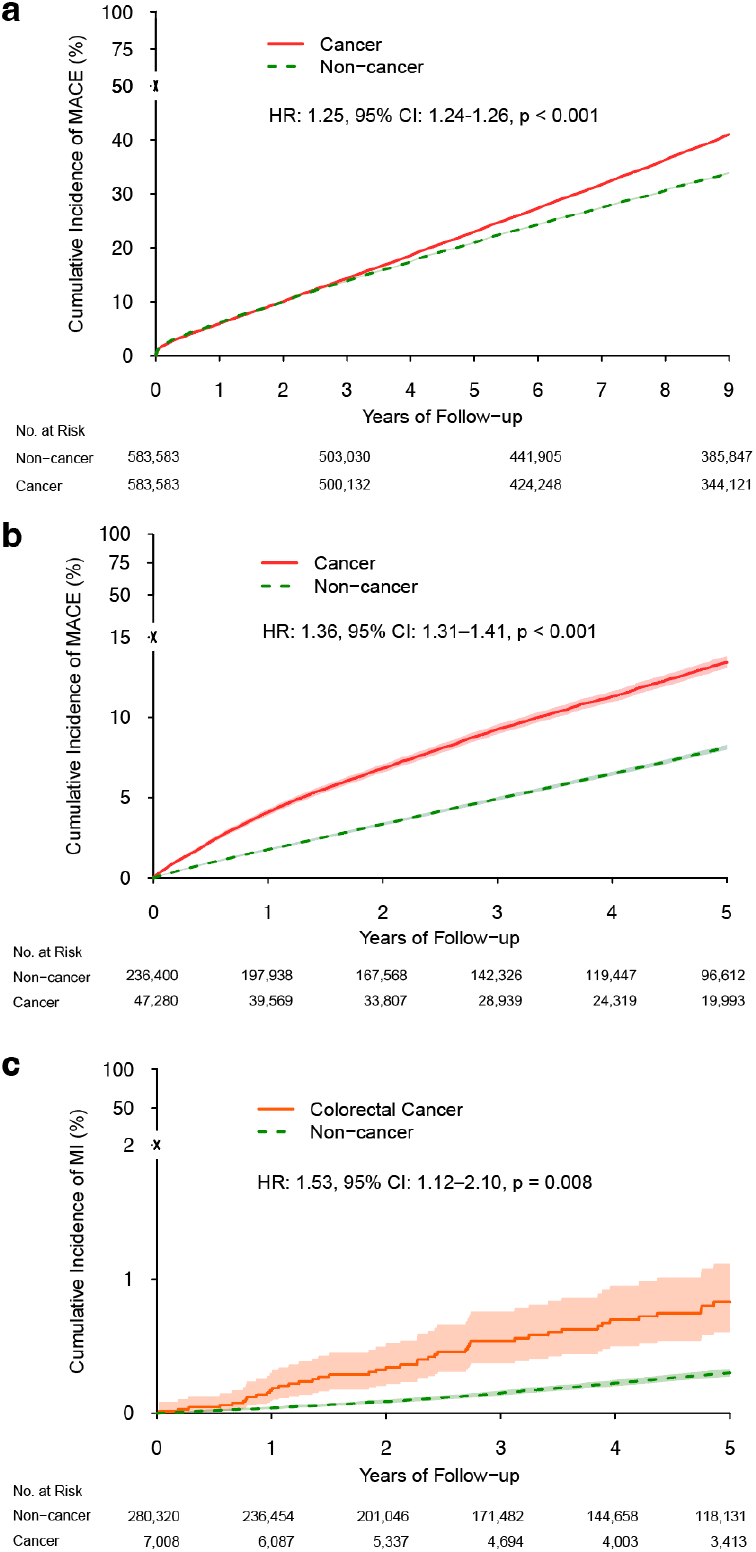
Cancer is associated with an increased risk of atherosclerotic cardiovascular events. **a**, Cumulative incidence of major adverse cardiovascular events (MACE) in patients with any cancer versus cancer-free individuals in the Allgemeine Ortskrankenkasse Bayern database. **b**, Cumulative incidence of MACE in patients with any cancer versus cancer-free controls in the Stanford Research Repository (STARR) cohort. **c**, Cumulative incidence of MI in colorectal cancer patients versus matched controls without colorectal cancer in the STARR cohort. For all panels, Hazard Ratios (HR), 95% confidence intervals (CI), and p-values were determined by Cox proportional hazards models.

### Impact of tumor implantation on plaque vulnerability

While the preceding observational data adjusted for available confounding factors, no retrospective analysis can fully exclude the existence of unmeasured residual sources of bias (e.g. changes in diet, physical activity, and/or intensity of disease surveillance and management following a cancer diagnosis) or account for potential late pro-atherosclerotic effects of anti-cancer therapies, even with rigorous propensity matching. To overcome these limitations, we next pursued a series of murine causation studies, using mouse models of atherosclerotic plaque rupture that had never been exposed to chemotherapy or any other pleiotropic risk factors such as smoking. Because colorectal carcinoma (CRC) is increasingly occurring in younger individuals(*23, 24*), has been linked to higher rates of CV mortality(*22*), and was the cancer most significantly associated with risk for MI in our epidemiology studies (Fig. 1c), this cancer was selected for our initial studies. We began by implanting syngeneic MC-38 colorectal cancer cells into the flanks of atheroprone *Apoe*^−/−^ mice fed a high-fat ‘Western’ diet. These mice began to develop larger atherosclerotic plaques (Fig. S2a,b), but the rapid rate of orthotopic tumor expansion limited our ability to study mice through the entire course of lesion development. In response, we pivoted to the ‘tandem stenosis’ (TS) model, which is a surgical model of plaque destabilization that manifests with accelerated atherosclerosis and carotid plaque rupture over a matter of weeks(*25*). In this accelerated model, the presence of a tumor was sufficient to drive a modest increase in lesion size over a shorter time course (Fig. 2a,b), but induced an even more striking increase in features of plaque vulnerability compared to animals without malignancy (Fig. 2c,d and Fig. S2c). Specifically, lesions in tumor-bearing mice demonstrated a marked increase in intraplaque endothelial staining, indicating enhanced neovessel ingrowth into the enlarging lesions. Furthermore, there was a significant increase in intraplaque hemorrhage, suggesting these nascent tubes were immature and prone to rupture(*26*). Subsequent Matrigel assays confirmed that secreted factors present in the supernatant of CRC cells were sufficient to augment angiogenesis *in vitro* (Fig. 2e), an effect that was not seen when endothelial cells were cultured with factors produced by healthy colonic epithelium. Together, these data provide evidence that the presence of a tumor, even distant from the atherogenic vascular bed, is sufficient to promote lesional neovessel formation and intraplaque hemorrhage. These effects promote lesion instability and plaque rupture and ultimately provide a potential explanation for the increased risk for MI observed in cancer survivors.

**Figure 2.**
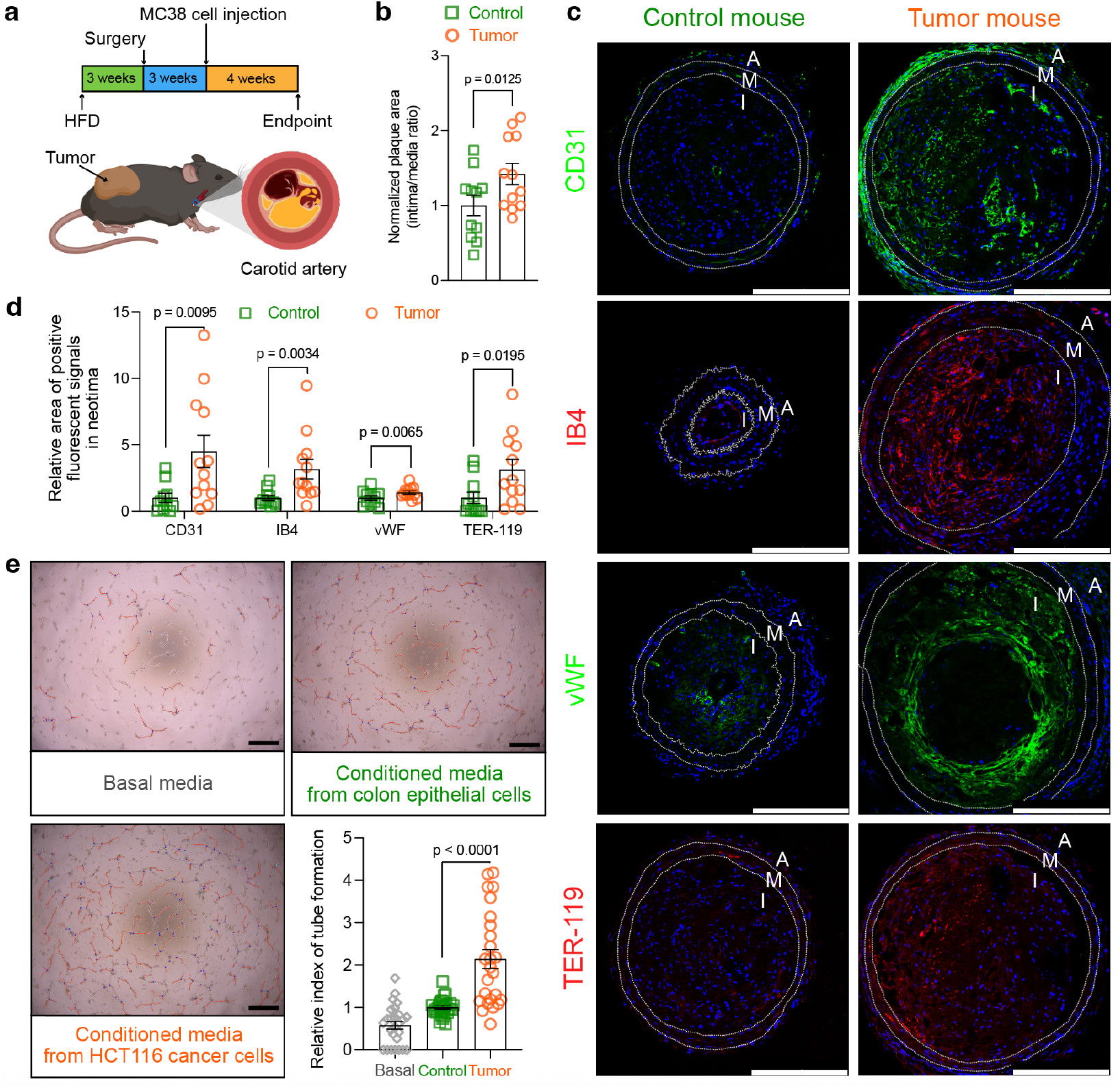
Colorectal cancer promotes intraplaque neovascularization and plaque vulnerability. **a**, Schematic of the animal model and experimental timeline. Atherosclerosis-prone *Apoe*−^/^− mice were fed a high-fat diet (HFD) for 10 weeks, with tandem stenosis of the carotid artery induced at 3 weeks and subcutaneous injection of MC38 colorectal cancer at 6 weeks. **b**, Quantification of plaque area (intima-to-media ratio) in carotid arteries from tumor-bearing (*n* = 12) and control (*n* = 11) mice, normalized to the control group. Data are presented as mean ± s.e.m. Statistical significance was assessed using a one-sample t-test. **c**, Representative immunofluorescence images of carotid artery sections stained for endothelial markers CD31, isolectin B4 (IB4), von Willebrand factor (vWF), and erythrocytes (TER-119). Scale bar, 200 μm. A, adventitia; M, media; I, intima. **d**, Quantification of the fluorescent signal area for the markers in c from tumor-bearing (*n* = 12) and control (*n* = 10-11) mice. Data are presented as mean ± s.e.m. Data are normalized to the control group and presented as mean ± s.e.m. Statistical significance was assessed using a one-sample t-test or Wilcoxon signed-rank test. **e**, Endothelial tube formation assay using human aortic endothelial cells (HAECs) treated with conditioned media from primary colonic epithelial cells (Control) or HCT116 colorectal cancer cells (Tumor). Representative images (top) and quantification (bottom), normalized to the control basal media group. Data are presented as mean ± s.e.m. *n* = 25-27 per group and are pooled from 5 independent experiments. Statistical significance was assessed using the Wilcoxon signed-rank test. Scale bar, 500 μm.

### Pro-inflammatory cytokines secreted by tumors drive angiogenic cascades in the atherosclerotic plaque

To investigate the mechanism by which cancer promotes plaque vulnerability, we next performed RNA sequencing on atherosclerotic aortic arch samples harvested from mice with and without tumors. Though dozens of genes are perturbed in the presence of a tumor (Fig. 3a), pathway analysis confirmed that the most differentially regulated pathway in the blood vessels of tumor-bearing mice was ‘angiogenesis’ (Fig. 3b). To refine and narrow the list of dysregulated genes, these RNA sequencing experiments were then replicated under a variety of conditions, including in young vs old mice and mice on a normal vs high-fat diet (Fig. 3c and Fig. S2d-h). This allowed us to identify a single gene that was consistently upregulated in the presence of colorectal cancer across all conditions, *Lrg1*. LRG1 is a relatively understudied glycoprotein that has previously been shown to promote pathological angiogenesis in response to TGFβ-dependent signaling and other disease-related stimuli(*27*). Single-cell analyses confirmed that *Lrg1* is most upregulated in vascular endothelial cells relative to other vascular cell types (Fig. 3d,e), and bulk RNA sequencing using several other tumor models confirmed consistent upregulation of vascular *Lrg1* across every cancer type tested, including breast (EO771), lung (LLC), and skin cancer (B16-F10) (Fig. S2i-l). This upregulation could be recapitulated in endothelial cells *in vitro* upon exposure to conditioned media collected from cultured CRC cells (Fig. 3f). Subsequent analysis of carotid endarterectomy (CEA) samples from patients undergoing surgery for symptomatic carotid disease (stroke, transient ischemic attack, or *amaurosis fugax*) or high-grade asymptomatic carotid stenosis confirmed a marked upregulation of intravascular LRG1 amongst cancer survivors, at both the mRNA and protein level, compared to cancer-free controls (Fig. 3g-i). To determine if these cancer-related changes translated into a pro-angiogenic effect within the human atheroma, we then conducted machine learning (ML)-guided analyses on an additional n=154 Elastica van Giesson (EvG)-stained CEA samples from the Munich Vascular Biobank. Using mutual information analysis, a history of a cancer diagnosis was among the top five predictors of intraplaque hemorrhage intensity, following only established risk factors of aging, diabetes, and smoking (Fig. 3j). Analysis of proteomic data from the UK Biobank confirmed an elevation of LRG1 in the circulation of cancer patients compared to those with no history of malignancy (Fig. 3k), and demonstrated that LRG1 expression directly correlated with risk for MACE events over the ensuing 5 years (Fig. S3a,b).

**Figure 3.**
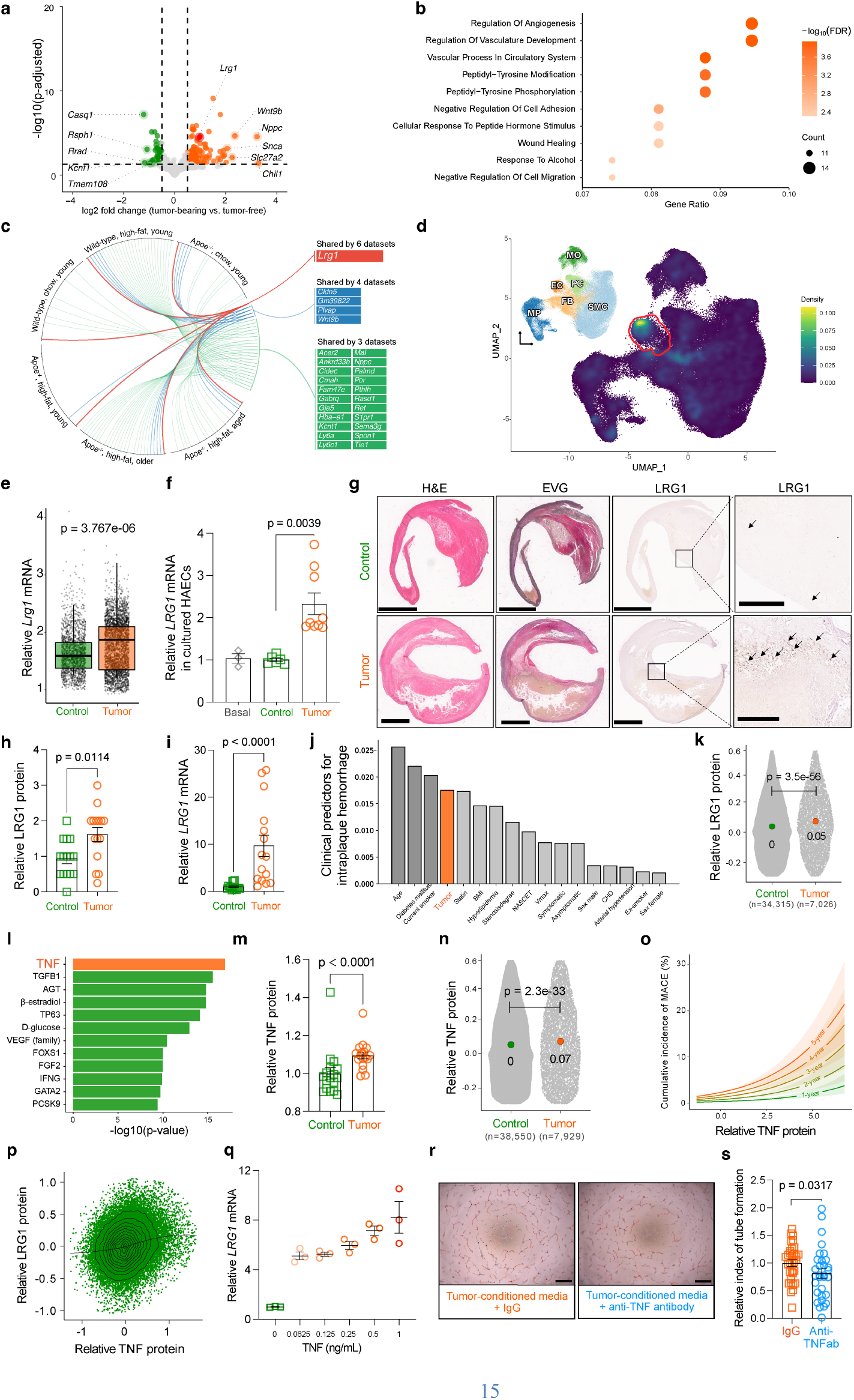
Tumors upregulate LRG1 expression in the atherosclerotic vasculature through TNF-α. **a**, Volcano plot of differentially expressed genes from bulk RNA sequencing of arteries from tumor-bearing versus control mice, with *Lrg1* highlighted. **b**, Gene Ontology (GO) enrichment analysis of upregulated genes showing enrichment for angiogenesis-related pathways. FDR, false discovery rate. **c**, Circos plot showing consistently upregulated genes, including *Lrg1*, across six different mouse atherosclerotic models. Variables were genotype (wild-type versus *Apoe*−^/^−), diet (chow diet versus HFD), and age (18 weeks to 1 year). **d**, UMAP visualization of single-cell RNA sequencing (scRNA-seq) data from arteries of tumor-bearing mice, showing the density of *Lrg1* expression by cell type. The endothelial cell cluster is outlined in red. **e**, Violin plot of *Lrg1* expression within endothelial cells from the scRNA-seq data from tumor-bearing and control mice (2 pooled samples from 4 mice per group). Boxes represent the interquartile range (IQR) with median; whiskers extend to 1.5× IQR. Statistical significance was assessed using the Mann-Whitney U test. **f**, Relative *Lrg1* mRNA expression in cultured HAECs treated with conditioned media from colonic epithelial cells or HCT116 tumor cells (*n* = 3-9 pooled from 3 independent experiments). Statistical significance was assessed using the Wilcoxon signed-rank test. **g**, Representative images of human carotid plaques stained with Hematoxylin and Eosin (H&E), Elastin-van Gieson (EVG), and immunostained for LRG1. Arrows indicate LRG1-positive microvessels. Scale bars: 2 mm (main), 400 μm (inset). **h, i**, Quantification of LRG1 protein (h) and mRNA (i) in carotid plaques from patients with cancer (n = 15) and matched cancer-free controls (n = 15). mRNA data were normalized to the control group. Data are presented as mean ± s.e.m. Statistical significance was assessed using the Mann-Whitney U test for protein and the Wilcoxon signed-rank test for mRNA. **j**, Feature importance of clinical predictors for intraplaque hemorrhage, with a history of cancer highlighted in orange. **k**, Violin plot of serum LRG1 protein levels in colorectal cancer patients (n = 7,026) versus matched controls (n = 34,315) from the UK Biobank. The y-axis shows *Olink* NPX values (log2 normalized protein expression). Statistical significance was assessed using the Mann-Whitney U test. **l**, Ingenuity Pathway Analysis (IPA) of arterial transcriptomics identifying tumor necrosis factor (TNF-α) as a top predicted upstream regulator. **m**, Relative serum TNF-α protein levels in tumor-bearing (n = 17) versus control (n = 17) mice revealed by *Olink*. Data are presented as mean ± s.e.m. Statistical significance was assessed using the Wilcoxon signed-rank test for mRNA. **n**, Serum TNF-α levels in colorectal cancer patients (n = 7,929) versus matched controls (n = 38,550) in the UK Biobank. Statistical significance was assessed using the Mann-Whitney U test. **o**, Modeled association between serum TNF-α levels and the cumulative incidence of MACE in the UK Biobank cohort over time horizons of one to five years. **p**, Scatter plot showing the positive correlation between serum LRG1 and TNF-α levels in the UK Biobank cohort (β = 0.159, *p* < 2.2e-16, R^2^ = 0.046). **q**, Relative *LRG1* mRNA expression in cultured HAECs treated with increasing concentrations of recombinant TNF-α (n = 3, experiments were replicated 3 times). **r, s**, Representative images (g) and quantification (h) of an endothelial cell tube formation assay using tumor-conditioned media pre-treated with a neutralizing anti-TNF-α antibody or an IgG isotype control. Data are presented as mean ± s.e.m. *n* = 31-32 per group and are pooled from 4 independent experiments. Statistical significance was assessed using a one-sample t-test. Scale bar, 500 μm.

To understand how this angiogenic factor is consistently upregulated in the vasculature of mice and humans with cancer, irrespective of cancer type or physical distance from the tumor, we next performed a series of upstream pathway analyses and unbiased proteomic studies on samples harvested from tumor-bearing mice. Ingenuity Pathway Analysis (IPA) software predicted that TNF-α was the upstream regulator most likely to drive the transcriptional changes observed in the plaques of cancerous mice (Fig. 3l). Direct measurement of circulating TNF-α levels confirmed a significant elevation in the bloodstream of mice with cancer (Fig. 3m), which was again observed in the circulation of cancer patients compared to cancer-free individuals using proteomic data from the UK Biobank (Fig. 3n and Fig. S3c). Additional analyses confirmed that TNF-α levels strongly correlated with risk for MACE(*28*) (Fig. 3o), possibly related to their positive relationship to LRG1 expression (Fig. 3p and Fig. S3d).

### Blockade of the TNF-α-LRG axis inhibits pathological angiogenesis and prevents cancer-dependent CVD

The preceding data suggest that TNF-α secreted by tumors may augment *Lrg1* expression, thus triggering an angiogenic process that accelerates intraplaque neovessel formation, lesional hemorrhage, and risk for plaque rupture. Given the clinical significance of this possibility, we next set out to determine its translational potential. First, we performed a series of cell culture studies to confirm that TNF-α directly induces *Lrg1* expression in endothelial cells (Fig. 3q) and that blockade of this axis could prevent tube formation *in vitro* (Fig. 3r,s).

Next, we proceeded to *in vivo* rescue studies, where mice implanted with colorectal cancer were subjected to carotid artery injury in the presence or absence of LRG1 knockdown (using shRNA) or TNF-α blockade (using an FDA-approved fusion protein, Etanercept) (Fig. 4a-d, and Fig. S3e). Compared to the pro-angiogenic effects of the tumors observed at baseline, intervention against the TNF-α-LRG axis reduced or completely eliminated the increase in endothelial tube ingrowth and intraplaque hemorrhage, even as the cancers continued to grow.

**Figure 4.**
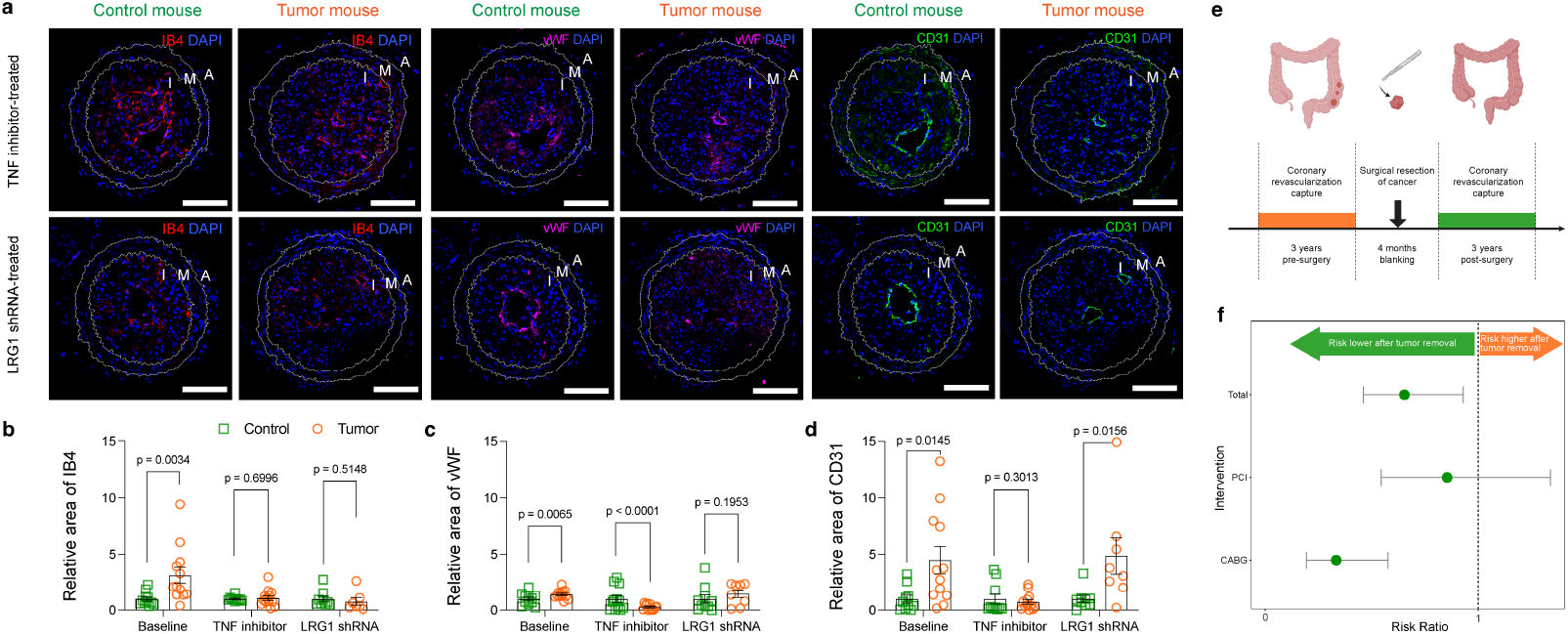
Blocking the TNF-LRG1 axis attenuates intraplaque angiogenesis and cardiovascular risk. **a**, Representative immunofluorescence images of carotid plaques from tumor-bearing *Apoe*−^/^− mice treated with a TNF-α inhibitor or LRG1-targeting shRNA (LRG1 shRNA), alongside their respective controls. Sections are stained for IB4, vWF, and CD31. Scale bar, 100 µm. **b-d**, Quantification of the relative immunostained area for vWF (b), IB4 (c), and CD31 (d) in plaques from the different treatment groups. Data are presented as mean ± s.e.m. *n* = 8-13 per group. Statistical significance was assessed using a one-sample t-test or the Wilcoxon signed-rank test. **e**, Schematic of the clinical study evaluating coronary revascularization incidence in patients before and after tumor resection surgery. **f**, Forest plot showing the relative risk of coronary revascularization procedures (percutaneous coronary intervention, PCI; or coronary artery bypass grafting, CABG) in the three years following tumor resection versus the three years preceding it. Total events (Ratio = 0.65; 95% CI, 0.46-0.93), PCI (Ratio = 0.85; 95% CI, 0.54-1.34), and CABG (Ratio = 0.33; 95% CI, 0.19-0.58). The analysis excludes a four-month perioperative period.

### Revascularization rates fall in the years following curative tumor resection

Finally, we attempted to explore the clinical relevance of these findings in humans by evaluating rates of coronary revascularization in the years before and after resection of early-stage colorectal tumors. This model was chosen given that stage I tumors can be surgically cured, without patients having to be exposed to concomitant radiation, chemotherapy, targeted therapy, or immunotherapy, each of which might otherwise confound the results. We utilized the Mass General Brigham Research Patient Data Registry to identify 6,920 patients with colon cancer who had undergone tumor resection alone without exposure to any of these potentially cardiotoxic therapies. We then compared their coronary revascularization event rates in the three years before surgery with the three years following surgery, with a 2-month blanking period before and after the surgical date to avoid the increased perioperative risk of cardiac event. Despite their increasing age, we found that the rates of coronary revascularization fell by 35% in the years following tumor removal (Fig. 4e,f). The effect was distinct from the pattern observed following prostate cancer resection (Fig. S4a,b), another tumor type that can be cured surgically, but has not been consistently linked to higher rates of CVD(*29, 30*).

## Discussion

Clinicians have long recognized the association between malignancy and a host of cardiovascular complications(*30-34*). These have classically been ascribed to the prothrombotic properties of cancer(*35*), the myopathic effects of anthracyclines(*32*), the pro-stenotic consequences of radiation therapy(*36*), and/or the pleiotropic harm derived from overlapping risk factors(*4, 30*). Recent reports have hinted at a more direct link between cancer and CVD; however, with large observational studies suggesting cardiovascular mortality rates up to twice as high in cancer survivors compared to the general population(*4, 22, 37*). Such observations have been bolstered by reports that more rigorously adjusted for cardiovascular co-morbidities and risk factors such as smoking, including findings from the Atherosclerosis Risk in Communities (ARIC) study(*3*) and those reported here. These data appear to be especially concerning for individuals diagnosed with cancer at a young age, who may have as much as a 10-fold increased risk of CAD(*31*), and CVD mortality rates that exceed the risk of death from the index cancer(*4*). Currently, approximately 5% of the US population are cancer survivors(*3*), but this statistic is predicted to nearly double by 2040(*4, 38*), due in part to the unexplained and dramatic increase in colorectal cancer diagnoses amongst younger individuals(*23*).

More than half a century ago, Folkman and colleagues described the role of angiogenesis in malignancy(*39*), and pathological neovascularization is now widely recognized as a key driver of oncogenesis. Intratumoral vessels not only perfuse the primary tumor, but also permit the escape of metastasizing cells into the circulation while maintaining the hemorrhagic, hypoxemic, and pro-oncogenic milieu of the tumor microenvironment(*40*). Similar phenomena occur in atherogenesis, where nascent tubes fail to recruit stabilizing pericytes, promoting the ingrowth of immature neovessels with reduced structural integrity(*26, 41, 42*). Because of their impaired barrier function, proatherogenic leukocytes are able to penetrate the developing plaque, and spontaneous hemorrhage from these fragile vessels can occur. These processes have been causally linked to plaque inflammation and lesion destabilization, and are considered drivers of plaque vulnerability and risk for clinical events. It is therefore interesting that while Lrg1 normally is viewed as an acute phase reactant typically restricted to the liver, it is also upregulated by inflammatory stimuli in both cancer and CVD, where it promotes the sprouting of vessels without mural cell coverage, enhances vascular permeability, and correlates with poor clinical outcomes(*27, 43, 44*). Thus, pathological Lrg1-dependent angiogenesis appears to be another mechanism which can be added to the growing list of shared biological processes known to drive both cancer and CVD, similar to chronic non-resolving inflammation, impaired efferocytosis, clonal hematopoiesis, and altered cell cycle regulation(*45*).

Importantly, this novel inter-disease cross-talk pathway appears to be targetable for translational purposes. Our data indicate that TNF-α secreted by tumors is both causally related to increased vascular Lrg1 expression, and associated with the heightened risk for adverse outcomes amongst cancer survivors. Numerous TNF-α inhibitors are FDA-approved for individuals with rheumatological conditions, and these have been associated with reduced vascular inflammation and potentially lower rates of MACE(*46, 47*). On the other hand, TNF-α inhibitors carry risks, including reactivation of certain latent malignancies(*48*) , and are contraindicated in moderate-to-severe heart failure, likely due to their effects on fluid retention. Thus, it is important that Lrg1 neutralizing antibodies have shown promising safety and efficacy in pre-clinical cancer and atherosclerosis models(*43, 44*), suggesting this axis may be a preferred target for clinical purposes, if the upstream driver itself is too risky to pursue.

Together, these results expand upon recent discoveries highlighting the bi-directional relationship between the world’s two leading killers. Cancer and CVD have long been known to share common risk factors, but recent work elegantly proved that CV disease also has the capacity to reprogram the immune system in such a way as to permit a tumorigenic environment and cancer growth, independent of hemodynamic changes(*19-21, 49*). Here we now show that the reverse is also true, and that cancer can have a direct causal effect on atherosclerosis, via an inflammatory and angiogenic pathway, that is independent of chemotherapy or smoking. These findings challenge conventional thinking about the compartmentalization of these two diseases and provide translational opportunities for the concerning increase of CVD amongst cancer survivors.

## Funding

National Institutes of Health grant R35HL 144475 (NJL)

National Institutes of Health grant R35HL 176060 (NJL)

American Heart Association Established Investigator Award EIA34770065 (NJL)

American Heart Association Postdoctoral Fellowship 23POST1012196 (LL)

American Heart Association Career Development Award 25CDA1452612 (LL)

Stanford Cancer Institute Innovation Award (NJL, LL)

Deutsche Gesellschaft für Innere Medizin (KUJ)

Else Kröner-Fresenius Stiftung 2024_EKES.02 (KUJ)

Corona-Stiftung S0199/10105/2024 (KUJ)

## Author contributions

Conceptualization: LL, RB, MVS, KTN, LM, HS, TGN, NJL

Methodology: LL, KUJ, RB, VHS, JH, MVS, DR, JK, NS, JLW, HW, HG, FW, AH, KTN, LM, HS, TGN, NJL

Investigation: LL, CF, RB, VHS, JH, MVS, DR, JK, NS, JLW, HW, HG, FW, SA, AH, MG.

Visualization: LL, VHS, JH, DR, JK, NS, JLW, HG, FW

Funding acquisition: LL, KUJ, NJL Supervision: NJL

Writing - original draft: LL, KUJ, NJL

Writing - review & editing: LL, CF, KUJ, RB, VHS, JH, MVS, DR, JK, NS, JLW, HW, HG, FW, KTN, LM, HS, TGN, NJL

## Data and materials availability

All sequencing data have been deposited in the Gene Expression Omnibus (GEO). Other data supporting the findings of this study are available in the main text, supplementary materials, or from the corresponding author upon request.

