## Supplementary materials for "Cancer drives atherosclerotic plaque vulnerability by inducing pathological angiogenesis"

**The PDF file includes:**

Materials and Methods
Figs. S1 to S4
Tables S1 to S10

### Materials and Methods

#### Human Cohort Studies

***Observational Registry (Allgemeine Ortskrankenkasse Bayern)***

To investigate the association between cancer and major adverse cardiovascular events (MACE), we analyzed German claims data from one of the country's largest statutory health insurers, covering approximately one-third of the national population (Allgemeine Ortskrankenkasse Bayern, AOK Bayern). This dataset, spanning January 2012 through December 2021, was used as the discovery cohort within the Observational Bavarian Health Insurance Registry, providing substantial statistical power.

Cohort selection is illustrated in **Fig. S1**. Diagnosis, procedure, and medication codes are listed in **Table S1**. Cancer patients and cancer-free controls were matched 1:1 using propensity scores based on age, sex, insulin use, statin use, and smoking status. Matching was performed without replacement, with a caliper width of 0.27 (two standard deviations of the logit of the propensity score). Balance was evaluated using standardized mean differences (defined as SMD < 0.1). Baseline covariates used in propensity score matching are provided in **Table S2**. Matching was implemented in Python (version 3.12.11) using the *scikit-learn* and *tqdm* packages.

Baseline characteristics after matching are summarized in **Tables S3 and S4**. Patients were followed from the index date until the occurrence of MACE, defined as all-cause mortality, ST-elevation myocardial infarction (STEMI, ICD-10 I21/I22), or ischemic stroke (I63/I64), or until administrative censoring on December 31, 2021.

Hazard ratios (HRs) and 95% confidence intervals (CIs) for MACE were estimated using Cox proportional hazards regression. Models were fitted in Python with the CoxPHFitter function from the *lifelines* package, which estimates HRs and 95% CIs via maximum partial likelihood, with p-values derived from Wald tests. Cumulative incidence functions (CIFs) were estimated in R (version 4.5.1) using the *tidycmprsk* package to visualize group differences in MACE incidence between cancer and non-cancer individuals.

***Stanford Research Repository (STARR)***

To validate findings from the German claims analysis and address limitations related to outpatient revascularization events and the treatment of death as a competing risk, we conducted a complementary validation analysis using the Stanford Research Repository (STARR) electronic health records (EHR) platform (release 2025_04_14). The data were harmonized to the OMOP Common Data Model.

Adults aged ≥18 years with at least 24 months of continuous observation were included. Cohort selection is illustrated in **Fig. S1**. Patients were categorized into cancer and non-cancer groups based on incident cancer diagnoses during a predefined run-in period. Individuals with a prior history of cancer or CVD were excluded. The primary endpoint was the occurrence of MACE within five years of follow-up.

Baseline covariates, including age, sex, smoking status, hyperlipidemia, hypertension, chronic kidney disease, obesity, and type 2 diabetes, were assessed before the index date. To account for baseline imbalances, 1:5 or 1:40 propensity score matching was applied. Baseline characteristics after matching are summarized in **Tables S5 and S6**. Competing risk regression analyses, including cause-specific Cox models (estimating the instantaneous rate of MACE among those still at risk) and Fine-Gray sub-distribution hazard models (estimating the cumulative incidence of MACE) as a comprehensive and rigorous statistical strategy, were performed to evaluate the association between cancer and MACE. Disease codes are provided in **Table S7**.

***UK Biobank***

*LRG1 or TNF levels from Olink Proteomics*

Proteomic data from 54,219 UK Biobank participants were analyzed to validate findings from unbiased murine proteomic studies and pathway analyses/GEO enrichment, including 7,968 individuals with a history of cancer, of whom 774 had colorectal cancer. Plasma LRG1 and TNF levels were measured using *Olink* proximity extension assays and stratified into quartiles. Associations between TNF levels and MACE were assessed using multivariable regression models adjusted for age, sex, and comorbidities. Disease codes are provided in **Table S8**.

***Munich Vascular Biobank***

*Histological analysis of human carotid artery lesions*

To assess differences in gene expression, with a focus on LRG1, human atherosclerotic carotid lesions from cancer patients and matched cancer-free patients were analysed. Cancer and non-cancer patients (n = 15 each) were matched for age and sex. All procedures involving human participants were approved by the Local Ethics Committee of the Technical University of Munich (approval no. 2799/10) and conducted in accordance with the Declaration of Helsinki. Written informed consent was obtained from all participants.

Carotid atherosclerotic plaques were obtained from patients undergoing carotid endarterectomy. Immediately after excision, tissue samples were transferred to pre-chilled RNAlater to preserve RNA integrity, and subsequently either fresh-frozen or processed for histological analysis. Samples were fixed in 4% paraformaldehyde (PFA) for 24 h. Decalcification was performed in EDTA-based solution (Entkalker soft SOLVAGREEN®, Carl ROTH, Karlsruhe, Germany) for 2-7 days. Specimens were then embedded in paraffin, and 2 μm sections were mounted on glass slides (Menzel SuperFrost, Fisher Scientific, Schwerte, Germany). Hematoxylin-eosin (HE) staining (ethanolic eosin Y solution, Mayer's acidic hemalum solution, Waldeck, Münster, Germany) and Elastica van Gieson (EvG) staining (picrofuchsin solution, Romeis 16th edition; Weigert's solution I, Romeis 15th edition) were carried out according to the manufacturers' protocols. Slides were mounted with Pertex (Histolab Products, Askim, Sweden) and glass coverslips (Engelbrecht, Edermünde, Germany).

LRG1 expression in human atherosclerotic plaques was assessed by immunohistochemistry (IHC) using a two-step biotin–streptavidin detection system (Bio SB) with an anti-LRG1 antibody (Invitrogen, PA5-81984). Staining was carried out according to the manufacturer's instructions. Counterstaining was performed with Mayer's hemalum solution (Carl Roth, Karlsruhe, Germany). Slides were scanned using an Aperio AT2 system (Leica, Wetzlar, Germany), and quantitative analysis of LRG1 staining was conducted with QuPath. Expression levels were compared between cancer and non-cancer groups (n = 15 per group).

*Image processing and hemorrhage mask generation*

EVG-stained histology images (RGB) of carotid plaques were processed to derive binary masks of intraplaque hemorrhage (IPH). Original images were down-scaled to accelerate processing and reduce noise. Color information was transformed from RGB to HSV space to enable hue-based separation of hemorrhage tones. K-means clustering was applied in HSV space to partition pixels into color clusters, and clusters corresponding to hemorrhage (orange-reddish tones in EVG) were filtered and merged. The resulting binary hemorrhage mask was up-scaled to the original image resolution for downstream quantification. This pipeline produced one "EVG-Image Hemorrhage-Mask" per slide and formed the basis for IPH quantifications.

*Clinical cohort and variables*

Analyses were conducted on an expanded cohort of n = 154 patients, including 21 samples from cancer patients and 133 samples of non-cancer patients. Clinical variables comprised sex, age, symptom status, affected side, hemorrhage (IPH, %), NASCET stenosis (%), an additional stenosis measure, Vmax (cm/s), arterial hypertension, hyperlipidemia, diabetes mellitus, smoking status, tumor status, coronary heart disease, BMI, and statin therapy. These features were curated from clinical and imaging records and were used in the mutual-information (MI) analysis below.

*Mutual-information analysis of clinical variables*

MI computations were performed in Python 3.9.19 (conda-forge) with NumPy 1.26.4, pandas 2.2.2, matplotlib 3.7.2, and scikit-learn 1.3.1.

Clinical data were read from Excel, with "x"/"X" entries recoded as missing (NaN) and all columns coerced to numeric. The target HEM (IPH, %) was discretized into three classes: (0–10] (class 1), (10–20] (class 2), and (>20) (class 3). Age was binned into four ranges: (0–55], (55–74], (74–84], (>84). Vmax (cm/s), NASCET stenosis percentage, Stenosis degree (%), and BMI were split into tertiles via a robust qcut implementation, with column names matched case-insensitively.

Discrete features (≤ 12 unique values or integer/categorical type) were imputed using the mode, continuous features using the median (SimpleImputer). MI between each feature and the 3-class HEM variable was computed with mutual_info_classification (random_state=0), results ranked, and displayed in a bar chart with tumor status highlighted.

***Mass General Brigham Research Patient Data Registry (RPDR)***

*Determining the effect of tumor resection on coronary revascularization*

To compare the incidence of coronary revascularization before and after tumor resection among cancer patients, data were extracted from the Mass General Brigham RPDR. Adult patients (≥18 years) diagnosed with colorectal or prostate cancer between 1990-01-01 and 2024-04-15 were included. Individuals with other malignancies or a history of prior cancer treatment were excluded. Patient selection within the RPDR was based on International Classification of Diseases version 9 and 10 (ICD-9, ICD-10) codes, and Current Procedural Terminology (CPT) codes. Demographics of patient cohorts are provided in **Table S9**. Diagnostic and procedural codes used for cohort identification and endpoint classification are listed in **Table S10**.

A two-month blanking period was applied before and after the date of surgical resection to minimize the potential confounding effects of perioperative events including pre-operative revascularization prior to cancer surgery and the risk of post-operative myocardial infarct. The pre-surgical observation window was defined as the three years preceding the start of the blanking period, while the post-surgical window comprised the three years following its conclusion.

Coronary revascularization events were defined using procedural codes for percutaneous coronary intervention (PCI) and coronary artery bypass grafting (CABG). The incidence of revascularization was compared between pre- and post-surgical periods. Given the nature of the experimental design, where each patient is their own control, there was no matching of patients pre- and post-surgery.

All analyses were performed using R (version 4.3.1). R package epitools was used to calculate risk ratios and p-values via Chi-squared.

**Mouse Models and Diet**

All animal experiments were approved by the Stanford University Administrative Panel on Laboratory Animal Care (APLAC protocol No. 27279) and conducted in accordance with the National Institutes of Health (NIH) guidelines for the care and use of laboratory animals. Apolipoprotein E-deficient (*Apoe*$^{-/-}$), wild-type C57BL/6J, or Myh11-Cre$^{ERT2}$, Rosa26$^{tdTomato/tdTomato}$, and *Apoe*$^{-/-}$ SMC-lineage-tracing mice were used in the study. Animals were either directly obtained from The Jackson Laboratory (Cat. Nos. 002052 and 000664) or bred in-house. Both sexes were utilized in the murine studies. Lineage-tracing studies were restricted to male mice, given that the reporter transgene is expressed on the y chromosome. Breast cancer studies were restricted to female mice. Mice were housed under specific pathogen-free conditions with ad libitum access to food and water. The animal facility was maintained on a 12-hour light/dark cycle at 22 °C. Mice were acclimated for 1-2 weeks before experimental procedures.

Beginning at 6 weeks of age, animals were fed either a standard chow diet or a high-fat diet (HFD) consisting of 21% anhydrous milk fat, 19% casein, and 0.15% cholesterol (Dyets Inc., Cat. No. 101511). In total, 170 mice were used across all experiments.

To investigate the effects of tumors on atherosclerosis progression, multiple mouse models were employed. In the classical atherosclerosis model, Myh11-Cre$^{ERT2}$, Rosa26$^{tdTomato/tdTomato}$, and *Apoe*$^{-/-}$ SMC-lineage-tracing mice were used. MC38 cells were inoculated subcutaneously at week 8 of HFD feeding, and mice were euthanized four weeks later.

In the tandem stenosis model, regular *Apoe*$^{-/-}$ mice were fed a high-fat diet (HFD) for 3 weeks, followed by tandem stenosis surgery on the right carotid artery via partial ligation at two consecutive sites with a 3 mm gap. Three weeks after surgery, $1 \times 10^4$ MC38 colon adenocarcinoma cells or Hanks' Balanced Salt Solution (HBSS) (Thermo Fisher Scientific, Cat. No. 14-175-095) were injected subcutaneously into both flanks to induce tumors. Mice were sacrificed four weeks after tumor inoculation under 2% Isospire™ (isoflurane) anesthesia.

For the multi-cancer comparison experiments, other tumor cell lines, including B16F10 melanoma, E0771 mammary adenocarcinoma, and Lewis lung carcinoma (LLC) cells, were used in place of MC38 cells.

For the anti-TNF treatment study, mice received subcutaneous injections of etanercept (Enbrel, 2 mg/kg; Pfizer) or isotype control (InVivoMAb™ recombinant human IgG1 Fc, Cat. No. BE0096) every other day, beginning two days prior to tumor implantation and continuing until sacrifice.

For vascular knockdown of *Lrg1*, wild-type mice were first injected intraperitoneally with AAV-m-PCSK9 ($1 \times 10^{11}$ genome copies; Vector Biolabs, Cat. No. AAV-268246) and placed on a high-fat diet (HFD) for two weeks. Subsequently, mice received intravenous injections of AAV-m-shLRG1 ($1.9 \times 10^{12}$ genome copies; Gene Universal Inc., Item No. V3001200-1) or AAV-scramble control. Tandem stenosis surgery was performed three weeks after AAV administration. The remaining procedures followed the standard tandem stenosis protocol as described above.

### Tissue Processing and Histological Analysis

Blood samples were collected via cardiac puncture and analyzed by the Stanford Animal Diagnostic Laboratory. Mice were then perfused with PBS (Corning, Cat. No. 21040CV), and tissues were dissected and either snap-frozen or fixed in 4% paraformaldehyde (Santa Cruz Biotechnology, Cat. No. sc-281692) overnight at 4 °C. Fixed tissues were subsequently incubated in 30% sucrose (Sigma-Aldrich, Cat. No. S0389-1KG) in PBS overnight at 4 °C until fully equilibrated. Samples were embedded in optimal cutting temperature (OCT) compound (Thermo Fisher Scientific, Cat. No. 23-730-571) and sectioned using a cryostat (Leica CM1950) at a thickness of 7 μm.

For atherosclerosis assessment, five sections spanning a total distance of 2 mm along the brachiocephalic artery (starting from the second carotid ligation site toward the aortic arch) were collected and analyzed per mouse. Intimal area was quantified using Oil Red O staining (Sigma-Aldrich, Cat. No. O1516-250ML) and defined as the area between the vessel lumen and the internal elastic lamina. Medial area was defined between the internal and external elastic laminae.

For immunofluorescence, cryosections were permeabilized with 0.1% Triton X-100 and blocked with Rodent Block M (Biocare Medical, Cat. No. RBM961H). Primary antibodies included anti-CD31 (Novus Biologicals, Cat. No. NB600-1475), anti-vWF (Abcam, Cat. No. ab287962), isolectin B4 (IB4; Vector Labs, Cat. No. B-1205-.5), and TER-119 (Santa Cruz, Cat. No. sc-19592). Sections were then incubated with appropriate secondary antibodies and counterstained with DAPI. Slides were mounted in Fluoroshield and imaged using a Leica Thunder Imager (QU-0117406-B). Vessel areas and fluorescence signal intensities were quantified in a blinded fashion using ImageJ.

LRG1 expression in human atherosclerotic plaques was assessed by immunohistochemistry (IHC) and quantitative reverse transcription PCR (RT-qPCR). Paraffin-embedded plaque sections were stained with an antibody against LRG1 and Elastica van Gieson to assess protein expression and location, as well as elastic lamina integrity. Staining intensity and mRNA levels were quantified and compared between cancer and non-cancer groups (n = 15 per group).

### RNA Extraction and Bulk RNA Sequencing

Aortic arches were homogenized in TRIzol reagent (Thermo Fisher Scientific, Cat. No. 15596018), and total RNA was extracted according to the manufacturer's instructions. RNA concentration and integrity were assessed using an Agilent 2100 Bioanalyzer. Sequencing libraries were generated using a poly(A) enrichment protocol and sequenced on an Illumina NovaSeq 6000 platform (Novogene, Sacramento, CA) to produce 150 bp paired-end reads.

Raw sequencing data were quality-checked using FastQC and trimmed using fastp. Clean reads were aligned to the mouse reference genome (GRCm38/mm10) using STAR (v2.7.5c) with default parameters. Gene-level quantification was performed using featureCounts (v2.0.3). Differential expression analysis was carried out using DESeq2 (v1.30.1) in R (v4.0.3), with significance defined as adjusted $p < 0.05$.

Pathway enrichment and upstream regulator analysis were performed using Ingenuity Pathway Analysis (IPA, QIAGEN).

### Single-cell RNA-seq Analysis

The aortic arch up to the level of the brachiocephalic artery was collected immediately after mouse euthanasia. Tissues were rinsed three times in PBS, minced, and enzymatically digested in HBSS with 2 U/mL Liberase (Sigma-Aldrich, Cat. No. 5401127001) and 2 U/mL elastase (Worthington, Cat. No. LS002279) at 37 °C for 1 hour. The resulting cell suspension was filtered through a 70 µm strainer, centrifuged at 500 × g for 5 minutes, and resuspended in fresh HBSS. Samples from three mice were pooled to generate each single-cell suspension.

Single-cell capture and library preparation were performed at the Stanford Functional Genomics Facility using the 10x Genomics Chromium platform, following the manufacturer's protocol. Libraries were prepared using the 10x Genomics single-cell 3' protocol and multiplexed for sequencing on the X Plus platform (Novogene), generating approximately 1.25 billion paired-end reads.

Sequencing data were processed using Seurat v4.4.0. Batch correction was performed using canonical correlation analysis (CCA). Gene expression normalization was conducted using SCTransform, which fits a Gamma-Poisson generalized linear model to account for sequencing depth and technical variation, producing log-transformed, normalized values. These were used for dimensionality reduction and clustering. Statistically significant principal components were identified via resampling-based significance testing and retained for Uniform Manifold Approximation and Projection (UMAP) visualization. Differential gene expression analysis across clusters was performed using a likelihood-ratio test. Genes were designated as cluster markers if they met the following criteria: expressed in ≥25% of cells in the cluster, adjusted $p < 0.05$, and log-fold change ≥ 0.25 compared to all other clusters.

**Quantitative PCR**

Total RNA was isolated from mouse tissues or cultured cells and reverse-transcribed into cDNA using the High-Capacity cDNA Reverse Transcription Kit (Thermo Scientific, Cat. No. 4368814). Quantitative PCR was performed using TaqMan Gene Expression Assays with probes for LRG1 (Hs00364835_m1, Thermo Scientific, Cat. No. 4331182) and GAPDH (Hs02786624_g1, Thermo Scientific, Cat. No. 4331182) or RPLP0 (HS00420895_gH, Thermo Fisher, Cat. No. 4331182) as house-keeping control.

**Tissue culture and cellular assays**

All cells were maintained in a humidified incubator at 37 °C with 5% $CO_2$. MC38 cells were cultured in DMEM (Corning, Cat. No. 10-013-CV) supplemented with 10% fetal bovine serum (FBS; Cytiva, Cat. No. SH30910.03) and 1% penicillin-streptomycin (P/S; Corning, Cat. No. 30-002-CI). Human aortic endothelial cells (HAECs) (ATCC, Cat. No. PCS-100-011) were cultured in EGM-2 endothelial cell growth medium (Lonza, Cat. No. CC-3162), and passages 2–5 were used for experiments. To assess HAEC response to tumor necrosis factor (TNF), cells were treated with recombinant human TNF (0.0625, 0.125, 0.25, 0.5, or 1 ng/mL; R&D Systems, Cat. No. 210-TA-005) for 4 hours prior to RNA extraction.

For experiments comparing conditioned media from primary versus cancer cells, HCT116 human colorectal cancer cells and primary human colon epithelial cells (Cell Biologics, Cat. No. H-6047) were used. HCT116 cells were maintained in McCoy's 5A medium (Thermo Fisher Scientific, Cat. No. 16600082), while primary epithelial cells were cultured in complete epithelial cell medium (Cell Biologics, Cat. No. H6621).

To block TNF signaling in the conditioned media, anti-TNF antibody (1 µg/µL; R&D Systems, Cat. No. MAB610-100) or an isotype control antibody (R&D Systems, Cat. No. MAB002) was added to the media prior to treatment.

For the tube formation assay, growth factor-reduced (GFR) Matrigel basement membrane matrix (Corning, Cat. No. 356230) was used to coat 48-well plates. HAECs ($2 \times 10^4$ cells/well) were seeded onto the coated wells. After 9 hours, tube structures were imaged, and quantification was performed using the Angiogenesis Analyzer plugin in ImageJ.

**Mouse Serum Proteomics**

Blood was collected at the endpoint of animal sacrifice (4 weeks after tumor cell inoculation), and serum was analyzed using the *Olink* Target 96 Mouse Exploratory Panel (Stanford Human Immune Monitoring Center). Differentially expressed proteins were identified and functionally annotated.

**Statistical Analysis**

Power calculations were performed a priori based on prior effect sizes and standard deviations. Data are presented as mean ± standard error of the mean (SEM). Statistical analyses were performed using GraphPad Prism 9. Outliers were identified and excluded using the ROUT method (Q = 1%). Normality was assessed with the Shapiro-Wilk test. For normally distributed data, parametric tests were applied (with Welch's correction when variances were unequal); otherwise, nonparametric tests were used. Comparisons between two groups were performed using an unpaired Student's t-test or a Mann-Whitney U test, as appropriate. For values normalized to the control group, a one-sample t-test or Wilcoxon signed-rank test was conducted.

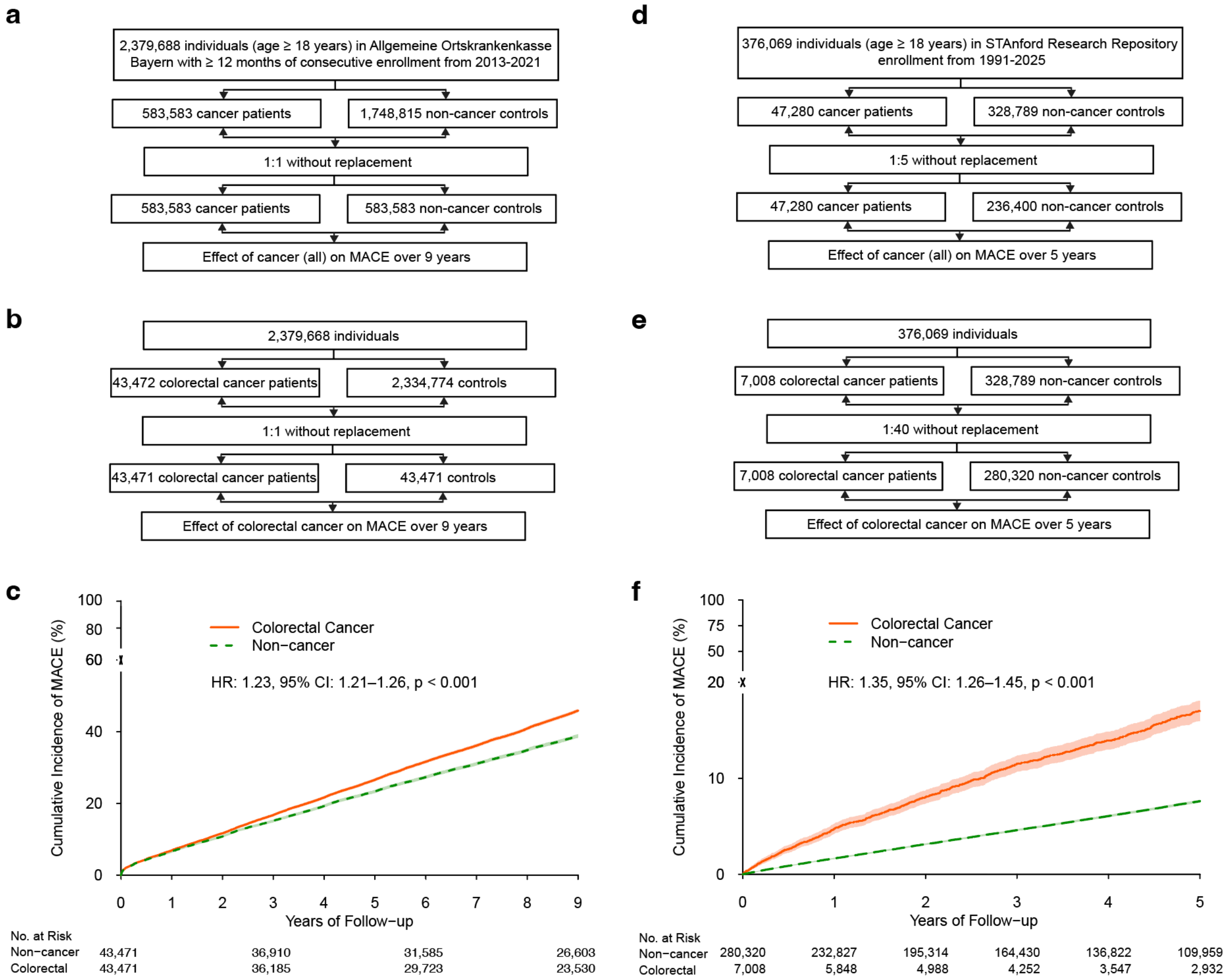


**Fig. S1. The clinical effect of cancer on major adverse cardiovascular events (MACE) in two independent population-based registries.**

**a**, Flowchart of cohort selection from the AOK Bayern registry for the all-cancer analysis. A total of 583,583 cancer patients were matched 1:1 with non-cancer controls and were followed for up to 9 years. The matching was based on age, sex, insulin use, statin use, and smoking status.

**b**, Flowchart of cohort selection from AOK Bayern for the colorectal cancer analysis.

**c**, Cumulative incidence of MACE in patients with colorectal cancer versus matched controls in AOK Bayern (HR = 1.23; 95% CI, 1.21-1.26; $p < 0.001$). The number of individuals at risk is indicated below the plot.

**d**, Flowchart of cohort selection from the STARR for the all-cancer analysis. A total of 47,280 cancer patients were matched 1:5 with non-cancer controls. The matching was based on age, sex, hyperlipidemia, hypertension, chronic kidney disease, obesity, diabetes, and smoking status.

**e**, Flowchart of cohort selection from STARR for the colorectal cancer analysis.

**f**, Cumulative incidence of MACE in patients with colorectal cancer versus matched controls in STARR (HR = 1.35; 95% CI, 1.26-1.45; $p < 0.001$). The number of individuals at risk is indicated below the plot.

For all panels, Hazard Ratios (HR), 95% confidence intervals (CI), and p-values were determined by Cox proportional hazards models.

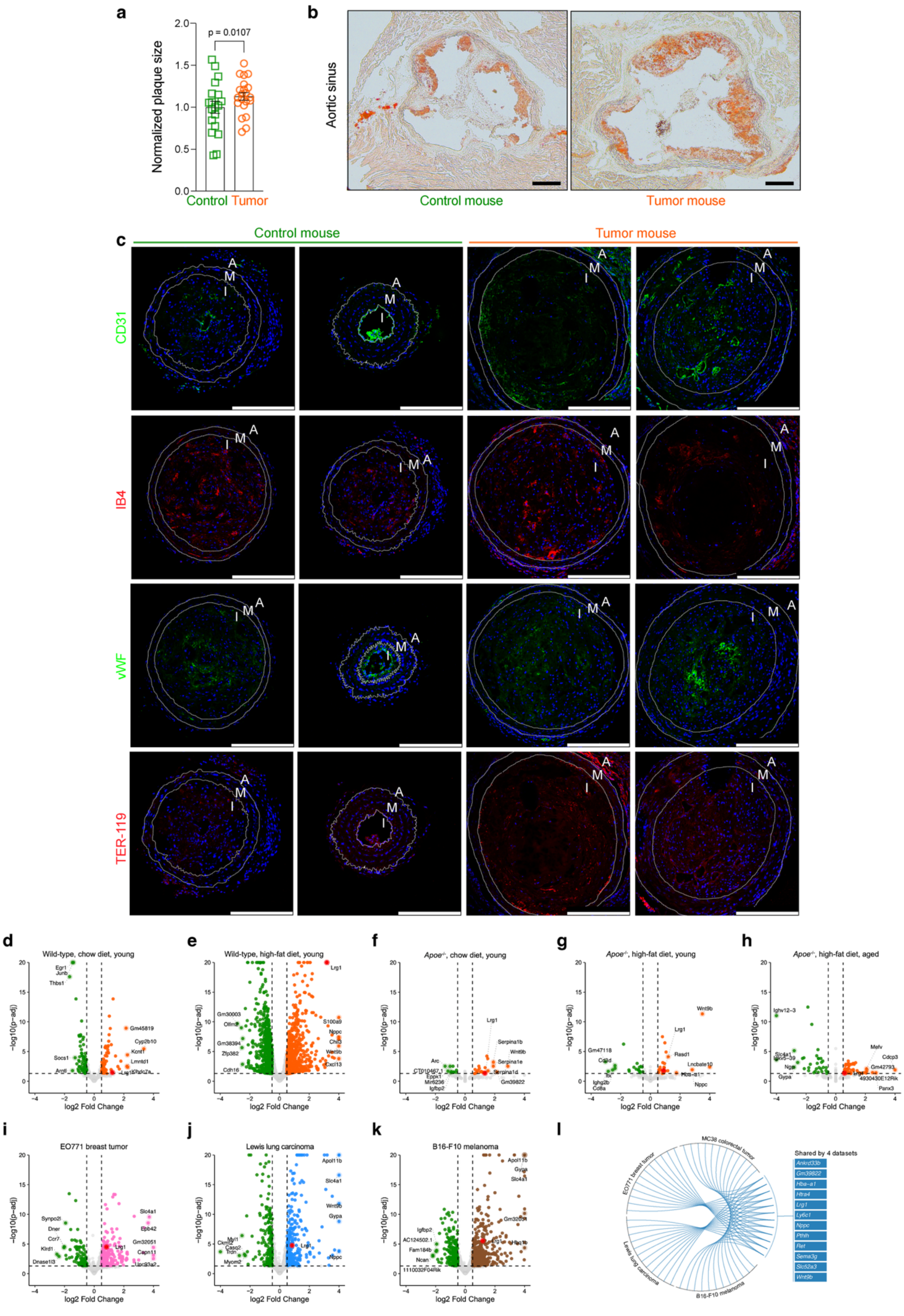
a
Normalized plaque size
p = 0.0107
Control
Tumor
b
Aortic sinus
Control mouse
Tumor mouse
c
Control mouse
Tumor mouse
CD31
IB4
vWF
TER-119
A
M
I
d
Wild-type, chow diet, young
e
Wild-type, high-fat diet, young
f
Apoe-/-, chow diet, young
g
Apoe-/-, high-fat diet, young
h
Apoe-/-, high-fat diet, aged
−log10(p−adj)
log2 Fold Change
i
EO771 breast tumor
j
Lewis lung carcinoma
k
B16-F10 melanoma
l
MC38 colorectal tumor
EO771 breast tumor
Lewis lung carcinoma
B16-F10 melanoma
Shared by 4 datasets
Ankrd33b
Gm39822
Hba-a1
Htra4
Lrg1
Ly6c1
Nppc
Pthlh
Ret
Sema3g
Slc52a3
Wnt9b

**Fig. S2. Tumor presence induces vascular remodeling and transcriptomic reprogramming across multiple models.**

**a**, Normalized plaque area in the aortic sinus of tumor-bearing mice (n = 21) versus tumor-free controls (n = 19), normalized to the tumor-free group. Data are presented as mean ± s.e.m. Statistical significance was assessed using a one-sample t-test.

**b**, Representative Oil Red O-stained sections of the aortic sinus from tumor-bearing and tumor-free mice, with insets showing plaque morphology. Scale bar, 200 µm.

**c**, Additional representative sections of the tandem stenosis segment from tumor-bearing and tumor-free mice. The sections are stained for CD31, IB4, vWF, and TER-119 to provide further evidence of tumor-induced vascular changes. Scale bar, 200 µm.

**d-k**, Volcano plots showing differentially expressed genes from bulk RNA sequencing of arteries from tumor-bearing versus control mice. The plots cover five datasets included in the Circos plot from Figure 3c as well as three additional tumor models (EO771 breast cancer, Lewis lung carcinoma [LLC], and B16-F10 melanoma).

**l**, Circos plot illustrating genes consistently upregulated in the arteries of *Apoe*$^{-/-}$ mice implanted with one of the four distinct tumor models: MC38 colorectal cancer, EO771 breast cancer, LLC, or B16-F10 melanoma. This panel highlights a core set of genes that may mediated the systemic effects of cancer on the vasculature.

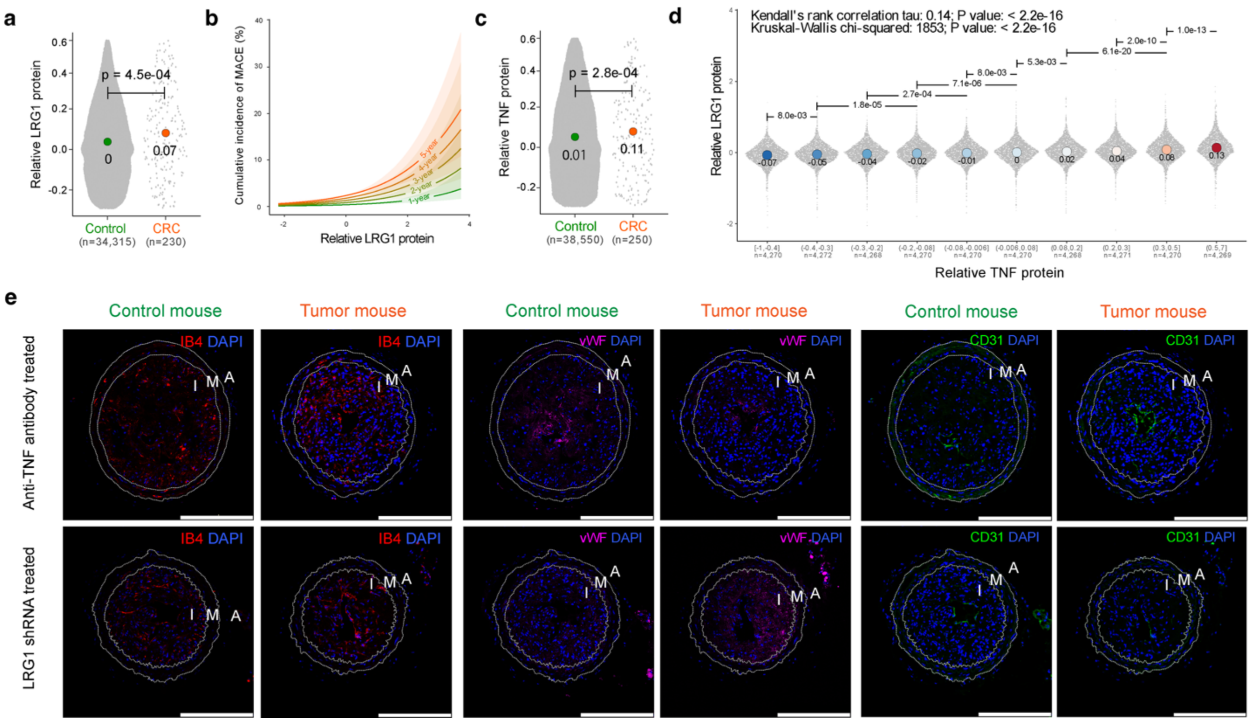


**Fig. S3. TNF-LRG1 signaling links cancer to cardiovascular risk and represents a therapeutic target.**

**a**, Serum levels of LRG1 in colorectal cancer patients compared to cancer-free controls from the UK Biobank (n = 34,315 controls, n = 230 colorectal cancer patients). The y-axis shows *Olink* NPX values (log2 normalized protein expression). Statistical significance was assessed using the Mann-Whitney U test.

**b**, Predicted cumulative incidence of MACE by serum LRG1 levels over different follow-up time windows (1 to 5 years) in the UK Biobank cohort.

**c**, Serum levels of TNF in colorectal cancer patients versus cancer-free controls from the UK Biobank (n = 38,550 controls, n = 250 colorectal cancer patients). The y-axis shows *Olink* NPX values (log2 normalized protein expression). Statistical significance was assessed using the Mann-Whitney U test.

**d**, Correlation between serum LRG1 and TNF levels in UK Biobank participants, demonstrating a positive association.

**e**, Additional representative histological images of carotid plaques from *Apoe*$^{-/-}$ mice with tumors. The top two row shows sections from mice treated with a TNF-neutralizing antibody versus an isotype control IgG. The bottom row shows sections from mice treated with AAV-shRNA targeting LRG1 versus a scramble control. Sections are stained for vWF, IB4, CD31, and DAPI to visualize blood vessels and cell nuclei. These images complement those in the main text and provide detailed morphological evidence of the therapeutic effect.

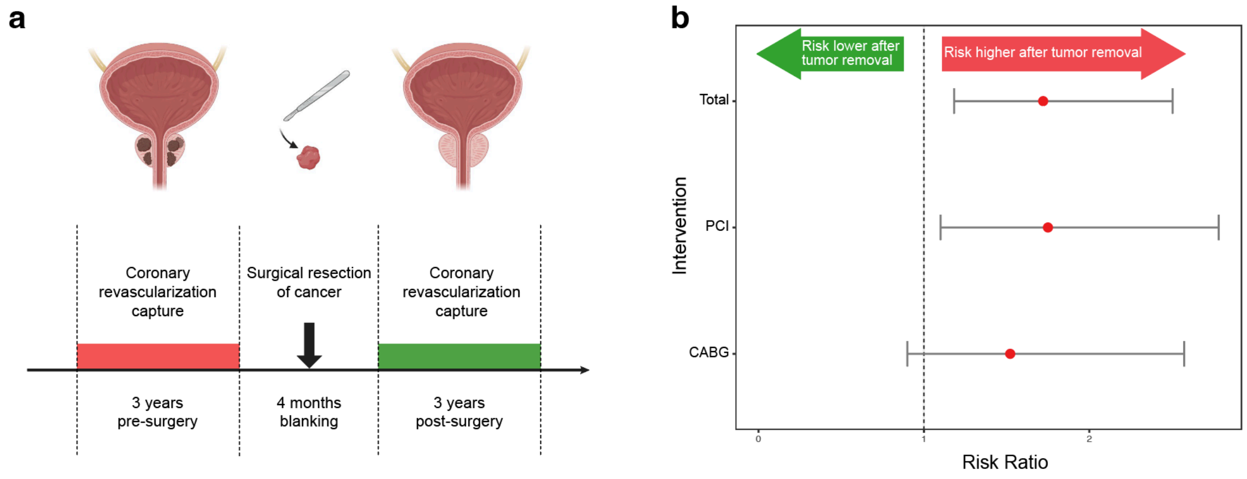


**Fig. S4. Clinical design and outcomes of a cohort evaluating coronary revascularization before and after tumor resection.**

**a**, Schematic illustrating the clinical design of the tumor resection cohort. The study evaluates the rate of coronary revascularization by PCI or CABG in the three-year period following surgical tumor resection compared to the three-year period preceding the surgery. The study includes a four-month blanking period immediately after surgery to exclude events directly related to the procedure.

**b**, Relative risk of coronary revascularization (PCI or CABG), with a relative risk of 1.0 indicating no change in risk. The plot shows the risk in the three years following versus preceding tumor resection for total events (Ratio = 1.72; 95% CI, 1.18–2.50), PCI (Ratio = 1.75; 95% CI, 1.10–2.78), and CABG (Ratio = 1.52; 95% CI, 0.90–2.57).

**Table S1. Diagnoses, procedures, and medication list. Provided are the International Classification of Diseases 10th Revision (ICD-10-GM), German procedure classification system (OPS), and Anatomical Therapeutic Chemical classification system (ATC) codes used in this study.**

| **Diagnosis / Procedure / Medication** | **Code** |
|---|---|
| ***International Classification of Diseases 10th Revision (ICD-10 GM)*** | |
| Malignant Neoplasms (*all for first screening*) | C00 – C97 |
| Diabetes Mellitus | E10, E11, E12, E13, E14 |
| Diabetes with End-organ Involvement | E11.2, E11.3, E11.4, E11.5, E10.2, E10.3, E10.4, E10.5 |
| Obesity | E66 |
| Dyslipidemia | E78 |
| Alcohol Abuse | F10 |
| Nicotine Abuse | F17 |
| Transient Ischemic Attack | G45 |
| Arterial Hypertension | I10, I15 |
| Malignant Hypertension | I10.1 |
| Hypertensive Heart and/or Kidney Disease | I11, I12, I13 |
| Unstable Angina Pectoris | I20.0 |
| Stable Angina Pectoris | I20.8 |
| Acute Myocardial Infarction | I21, I22 |
| ST- Segment Elevation Myocardial Infarction | I21.0, I21.1, I21.2, I21.3, I22.0, I22.1, I22.8 |
| Non-ST Elevation Myocardial Infarction | I21.4 |
| Stent Thrombosis | I25.16 |
| Chronic Ischemic Heart Disease | I25 |
| Chronic Ischemic Heart Disease with at least 2 vessel disease | I25.12, I25.13 |
| Atherosclerotic Heart Disease | I25.1 |
| Old Myocardial Infarction | I25.2 |
| Pulmonary Embolism | I26 |
| AV Block Grade II and III | I44.1, I44.2 |
| Other Cardiac Arrhythmias | I49 |
| Atrial Fibrilation and Flutter | I48 |
| Sick Sinus Syndrome | I49.5 |
| Chronic Heart Failure | I50 |
| Intracranial Hemorrhage | I61 |
| Bleeding | I60, I61, I62, K92.0, K92.1, K92.2 |
| Acute Stroke | I63, I64 |
| Intracranial Arteriovenous Malformation or Intracranial Aneurysm | I67.1 |
| Atherosclerosis | I70 |
| Aortic Atherosclerosis | I70.0 |
| Renal Atherosclerosis | I70.1 |
| Carotid stenosis | I65.2 |

| | |
|---|---|
| Peripheral Artery Disease | I70.2 |
| Arterial Embolism Or Thrombosis | I74 |
| Venous Thrombosis | I80, I81, I82 |
| Pneumonia | J13, J14, J15, J16 |
| COPD | J44 |
| Asthma | J45 |
| Acute Kidney Failure | N17 |
| Chronic Kidney Disease | N18 |
| Chronic Kidney Failure Stage 1 And 2 | N18.1, N18.2 |
| Chronic Kidney Failure Stage 3, 4 And 5 | N18.3, N18.4, N18.5 |
| Chronic Renal Insufficiency Requiring Dialysis | N18.5 |
| Other Chronic Kidney Disease | N18.8, N18.9 |

***German procedure classification system (OPS)***

| | |
|---|---|
| Diagnostic Cardiac Catheterization | 1-275 |
| (Percutaneous) Transluminal Stenting | 8-837 (excl.: 8-837.7, 8-837.8, 8-837.9, 8-837.a, 8-837.b, 8-837.c, 8-837.d, 8-837.e, 8-837.f, 8-837.g, 8-837.h, 8-837.j, 8-837.s) |
| Coronary Artery Bypass Graft Surgery | 5-36 |
| Complex treatment | 8-98 |

***Anatomical Therapeutic Chemical classification system (ATC)***

| | |
|---|---|
| Heparin | B01AB |
| Antiarrhythmics | C01B, C01AA |
| Nitrates | C01DA |
| Antihypertensives | C02 |
| Thiazides | C03A, C03EA, C07B, C07D, C09BA21, C09BA22, C09BA23, C09BA25, C09BA26, C09BA27, C09BA28, C09BA29, C09BA33, C09BA35, C09BA54, C09DA21, C09DA22, C09DA23, C09DA24, C09DA26, C09DA27, C09DA28 |
| Other Diuretics | C03B, C03DB, C03EA, C03EB, C03EC, C03X, C09BA01, C09BA02, C09BA03, C09BA04, C09BA05, C09BA06, C09BA07, C09BA08, C09BA09, C09BA12, C09BA13, C09BA15, C09DA01, C09DA02, C09DA03, C09DA04, C09DA06, C09DA07, C09DA08, C09DA09, C09DA10 |
| Loop Diuretics | C03C, C03EB, C03ED, C07C, C07D, C09BA55 |
| Aldosteron Antagonists | C03DA, C03EC, C03ED |

| | |
|---|---|
| Beta Blocking Agents | C07A, C07B, C07C, C07D, C07E, C07FB |
| Calcium Channel Blockers | C07FB, C08, C09BB, C09DB |
| ACE Inhibitors | C09A, C09BA01, C09BA02, C09BA03, C09BA04, C09BA05, C09BA06, C09BA07, C09BA08, C09BA09, C09BA12, C09BA13, C09BA15, C09BA21, C09BA22, C09BA23, C09BA25, C09BA26, C09BA27, C09BA28, C09BA29, C09BA33, C09BA35, C09BA54, C09BA55, C09BB |
| Angiotensin II Receptor Blockers | C09CA, C09DA01, C09DA02, C09DA03, C09DA04, C09DA06, C09DA07, C09DA08, C09DA09, C09DA10, C09DA21, C09DA22, C09DA23, C09DA24, C09DA26, C09DA27, C09DA28, C09DB |
| Angiogenesis Inhibitors | C09DX |
| Statins | C10AA |
| Other Lipid Modifying Agents | C10AB, C10AC, C10AD, C10AX |
| Oral Immunosuppressants | L04 |
| NSAIDs | M01A, M01BA, N02BA |
| Opioids | N02A |
| Non-Opioid Analgesics | N02BB, N02BE, N02BG |
| Drugs for obstructive airway diseases | R03 |

**Table S2**. **Baseline pre-exposure variables for propensity score matching.** Provided are the individual variables for the applied baseline propensity score matching used in this study.

| | |
|---|---|
| 1 | Age |
| 2 | Sex |
| 3 | Nicotine Abuse |
| 4 | Insulin intake |
| 5 | Statin intake |

**Table S3. Demographics of Propensity-matched Cohort (for all cancers)**

| Characteristic | All cancer | No cancer | p-value* | Standardized mean difference (SMD) |
|---|---|---|---|---|
| **Overall** | 583,583 | 583,583 | | |
| **Age Category** | | | 0.071 | 0.004 |
| 18-39 | 11,640 (2%) | 11,640 (2%) | | |
| 40-49 | 35,208 (6%) | 35,208 (6.1%) | | |
| 50-59 | 88,728 (15.2%) | 88,728 (15.2%) | | |
| 60-64 | 64,378 (11%) | 64,378 (11%) | | |
| 65-69 | 68,371 (11.7%) | 68,562 (11.8%) | | |
| 70-79 | 198,107 (34%) | 196,394 (33.7%) | | |
| 80+ | 116,869 (20%) | 118,391 (20.3%) | | |
| **Sex** | | | <0.001 | 0.010 |
| Male | 281,614 (48.3%) | 278,609 (47.7%) | | |
| Female | 301,969 (51.7%) | 304,974 (52.3%) | | |
| **Diabetes** | | | <0.001 | 0.011 |
| No | 430,634 (73.8%) | 433,328 (74.3%) | | |
| Yes | 152,949 (26.2%) | 150,255 (25.7%) | | |
| **Hypertension** | | | <0.001 | 0.071 |
| No | 175,904 (30.1%) | 195,277 (33.5%) | | |
| Yes | 407,679 (69.9%) | 388,306 (66.5%) | | |
| **Smoking?** | | | 0.651 | -0.001 |
| No | 553,625 (94.9%) | 553,516 (94.8%) | | |
| Yes | 29,958 (5.1%) | 30,067 (5.2%) | | |
| **Insulin use?** | | | <0.001 | -0.010 |
| No | 553,640 (94.9%) | 552,376 (94.7%) | | |
| Yes | 29,943 (5.1%) | 31,207 (5.3%) | | |
| **Obesity** | | | <0.001 | 0.028 |
| No | 485,939 (83.3%) | 491,935 (84.3%) | | |
| Yes | 97,644 (16.7%) | 91,648 (15.7%) | | |
| **Chronic Kidney Disease** | | | <0.001 | 0.034 |
| No | 529,697 (90.8%) | 535,309 (91.7%) | | |
| Yes | 53,886 (9.2%) | 48,274 (8.3%) | | |
| **Hyperlipidemia** | | | <0.001 | 0.047 |
| No | 342,713 (58.7%) | 356,099 (61%) | | |
| Yes | 240,870 (41.3%) | 227,484 (39%) | | |
| **Old MI** | | | 0.001 | -0.006 |
| No | 563,680 (96.6%) | 563,020 (96.5%) | | |
| Yes | 19,903 (3.4%) | 20,563 (3.5%) | | |
| **Congestive Heart Failure** | | | 0.046 | -0.004 |
| No | 502,847 (86.2%) | 502,101 (86%) | | |

| | | | | |
|---|---|---|---|---|
| Yes | 80,736 (13.8%) | 81,482 (14%) | | |
| **Peripheral Artery Disease** | | | 0.005 | 0.009 |
| No | 562,832 (96.4%) | 563,351 (96.5%) | | |
| Yes | 20,751 (3.6%) | 20,232 (3.5%) | | |
| **Statin Use?** | | | 0.943 | 0.000 |
| No | 445,224 (76.3%) | 445,258 (76.3%) | | |
| Yes | 138,359 (23.7%) | 138,325 (23.7%) | | |

**Table S4. Demographics of Propensity-matched Cohort (colorectal cancer)**

| Characteristic | Colorectal cancer | No cancer | p-value* | Standardized mean difference (SMD) |
|---|---|---|---|---|
| **Overall** | 43,471 | 43,471 | | |
| **Age Category** | | | 1.00 | 0.000 |
| 18-39 | 417 (0.9%) | 417 (0.9%) | | |
| 40-49 | 1,892 (4.4%) | 1,892 (4.4%) | | |
| 50-59 | 5,976 (13.7%) | 5,976 (13.7%) | | |
| 60-64 | 4,429 (10.2%) | 4,429 (10.2%) | | |
| 65-69 | 5,090 (11.7%) | 5,090 (11.7%) | | |
| 70-79 | 15,538 (35.8%) | 15,538 (35.8%) | | |
| 80+ | 10,120 (23.3%) | 10,120 (23.3%) | | |
| **Sex** | | | 1.00 | 0.000 |
| Male | 23,270 (53.5%) | 23,270 (53.5%) | | |
| Female | 20,201 (46.5%) | 20,201 (46.5%) | | |
| **Diabetes** | | | <0.001 | 0.039 |
| No | 30,895 (71.1%) | 31,647 (72.8%) | | |
| Yes | 12,576 (28.9%) | 11,824 (27.2%) | | |
| **Hypertension** | | | <0.001 | 0.035 |
| No | 12,901 (29.7%) | 13,607 (31.3%) | | |
| Yes | 30,570 (70.3%) | 29,864 (68.7%) | | |
| **Smoking?** | | | 1.00 | 0.000 |
| No | 41,637 (95.8%) | 41,637 (95.8%) | | |
| Yes | 1,834 (4.2%) | 1,834 (4.2%) | | |
| **Insulin use?** | | | 1.00 | 0.000 |
| No | 40,828 (93.9%) | 40,828 (93.9%) | | |
| Yes | 2,643 (6.1%) | 2,643 (6.1%) | | |
| **Obesity** | | | <0.001 | 0.036 |
| No | 36,066 (83%) | 36,645 (84.3%) | | |
| Yes | 7,405 (17%) | 6,826 (15.7%) | | |
| **Chronic Kidney Disease** | | | 0.002 | 0.022 |
| No | 39,099 (89.9%) | 39,377 (90.6%) | | |
| Yes | 4,372 (10.1%) | 4,094 (9.4%) | | |
| **Hyperlipidemia** | | | 0.539 | 0.004 |
| No | 25,800 (59.3%) | 25,890 (59.6%) | | |
| Yes | 17,671 (40.7%) | 17,581 (40.4%) | | |
| **Old MI** | | | 0.783 | -0.002 |
| No | 41,934 (96.5%) | 41,918 (96.4%) | | |
| Yes | 1,537 (3.5%) | 1,553 (3.6%) | | |
| **Congestive Heart Failure** | | | 0.069 | 0.012 |
| No | 36,784 (84.6%) | 36,977 (85.1%) | | |

| | | | | |
|---|---|---|---|---|
| Yes | 6,687 (15.4%) | 6,494 (14.9%) | | |
| **Peripheral Artery Disease** | | | 1.000 | -0.000 |
| No | 41,876 (96.3%) | 41,875 (96.3%) | | |
| Yes | 1,595 (3.7%) | 1,596 (3.7%) | | |
| **Statin Use** | | | 1.000 | 0.000 |
| No | 33,491 (77%) | 33,491 (77%) | | |
| Yes | 9,980 (23%) | 9,980 (23%) | | |

**Table S5. Demographics of matched Cohort (all cancers)**

| Characteristic | Overall<br>N = 283,680[1] | non-cancer<br>N = 236,400[1] | cancer<br>N = 47,280[1] | p-value[2] |
|---|---|---|---|---|
| Gender | 165,575 (58%) | 138,457 (59%) | 27,118 (57%) | <0.001 |
| Age | 45 (35, 59) | 43 (34, 57) | 54 (40, 66) | <0.001 |
| Hyperlipidemia | 35,932 (13%) | 28,740 (12%) | 7,192 (15%) | <0.001 |
| Hypertension | 38,781 (14%) | 29,713 (13%) | 9,068 (19%) | <0.001 |
| CKD | 4,175 (1.5%) | 3,232 (1.4%) | 943 (2.0%) | <0.001 |
| Obesity | 16,965 (6.0%) | 13,770 (5.8%) | 3,195 (6.8%) | <0.001 |
| T2DM | 14,751 (5.2%) | 11,447 (4.8%) | 3,304 (7.0%) | <0.001 |
| Smoking | 58,159 (21%) | 45,500 (19%) | 12,659 (27%) | <0.001 |

[1] n (%); Median (Q1, Q3)
[2] Pearson's Chi-squared test; Wilcoxon rank sum test

**Table S6. Demographics of matched Cohort (colorectal cancer)**

| Characteristic | Overall N = 287,328[1] | non-cancer N = 280,320[1] | cancer N = 7,008[1] | p-value[2] |
|---|---|---|---|---|
| Gender | 140,938 (49%) | 137,959 (49%) | 2,979 (43%) | <0.001 |
| Age | 40 (31, 55) | 39 (31, 54) | 58 (52, 67) | <0.001 |
| Hyperlipidemia | 34,018 (12%) | 32,210 (11%) | 1,808 (26%) | <0.001 |
| Hypertension | 32,675 (11%) | 30,710 (11%) | 1,965 (28%) | <0.001 |
| CKD | 3,618 (1.3%) | 3,435 (1.2%) | 183 (2.6%) | <0.001 |
| Obesity | 15,989 (5.6%) | 15,434 (5.5%) | 555 (7.9%) | <0.001 |
| T2DM | 12,686 (4.4%) | 11,954 (4.3%) | 732 (10%) | <0.001 |
| Smoking | 47,625 (17%) | 45,516 (16%) | 2,109 (30%) | <0.001 |

[1] n (%); Median (Q1, Q3)
[2] Pearson's Chi-squared test; Wilcoxon rank sum test

**Table S7. ICD codes for STARR**

| Conditions | Diagnosis or procedure codes | |
|---|---|---|
| Cardiovascular disease (CVD) | Coronary artery disease | *Diagnosis*:<br>(ICD-9) 410.xx (acute myocardial infarction); 411.xx (other acute and subacute forms of ischemic heart disease); 412 (old myocardial infarction); 414.0x, 414.2 - 414.9 (coronary atherosclerosis); 429.7x (sequelae of myocardial infarction)<br>(ICD-10) I21.x (acute myocardial infarction; excluding I21.Ax); I22.x (STEMI/NSTEMI); I23.x (complications following STEMI/NSTEMI); I24.x (other acute ischemic heart disease); I25.x (chronic ischemic heart disease; excluding I25.3, I25.4x); Z95.5 (presence of coronary angioplasty implant and graft); Z98.61 (coronary angioplasty status).<br>T82.211, T82.211A, T82.211D, T82.211S, T82.212, T82.212A, T82.212D, T82.212S, T82.213, T82.213A, T82.213D, T82.213S, T82.218, T82.218A, T82.218D, T82.218S, Z951<br>*Procedure*:<br>(ICD-9 CM) 0.66 (PCTA); 36.0x (removal of coronary artery obstruction); 36.1x (bypass anastomosis for heart revascularization); 36.2 (heart revascularization by arterial implants); 36.3x (other heart revascularization); 17.55 (transluminal coronary atherectomy).<br>(ICD-10 PCS Diagnosis) 0210xxx (coronary bypass, one artery); 0211xxx (coronary bypass, two arteries); 0212xxx (coronary bypass, three arteries); 0213xxx (coronary bypass, four arteries); 0270xxx (coronary dilation, one artery); 0271xxx (coronary dilation, two arteries); 0272xxx (coronary dilation, three arteries); 0273xxx (coronary dilation, four arteries)<br>(CABG) B2020ZZ, B2021ZZ, B202YZZ, B2030ZZ, B2031ZZ, B203YZZ, B212010, B2120ZZ, B212110, B2121ZZ, B212Y10, B212YZZ, B213010, B2130ZZ, B213110, B2131ZZ, B213Y10, B213YZZ, B22300Z, B2230ZZ, B22310Z, B2231ZZ, B223Y0Z, B223YZZ, B223Z2Z, B223ZZZ, B233Y0Z, B233YZZ, B233ZZZ<br>(CPT) (PCI) 92920, 92921, 92924, 92925, 92928, 92929, 92933, 92934, 92937, 92938, 92941, 92943, 92944, C9600, C9601, C9602, C9603, C9604, C9605, C9606, C9607, C9608<br>(CABG) 33510, 33511, 33512, 33513, 33514, 33516, 33517, 33518, 33519, 33520, 33521, 33522, 33523, 33525, 33528, 33530, 33533, 33534, 33535, 33536, |

| | | |
|---|---|---|
| | | 35600, 4110F, 75762, 75764, 75766, 75767, 93551, C9604, C9605, G8158, G8159, G8160, G8161, G8162, G8163, G8164, G8165, G8166, G8167, G8170, G8171, G8172, G8497, G8544, G8573, G8574 |
| | Peripheral artery disease, aortic atherosclerosis | *Diagnosis*:<br>(ICD-9) 433.xx (occlusion and stenosis of precerebral arteries); 440.xx (atherosclerosis); 443.9 (peripheral vascular disease, unspecified)<br>(ICD-10) I65.xx (occlusion and stenosis of precerebral arteries, not resulting in cerebral infarction); I70.xxx (atherosclerosis); I73.9 (peripheral vascular disease, unspecified); I75.xxx (atheroembolism)<br>*Procedure*:<br>Peripheral Revascularization (carotid and cerebral included):<br>(ICD-9 CM) 00.55 (insertion of drug eluting non-coronary stent); 00.63 (perc insertion of carotid artery stent); 00.64 (perc insertion of extracranial artery stent); 00.65 (perc insertion of intracranial artery stent); 17.53 (perc atherectomy extracranial vessel); 17.54 (perc atherectomy intracranial vessel); 17.56 (atherectomy of noncoronary vessel); 38.1x (endarterectomy); 39.25, 39.26, 39.29 (peripheral bypass grafting); 39.50 (Angioplasty of other non-coronary vessel(s)); 39.90 (Insertion of non- drug-eluting peripheral (non-coronary) vessel stent(s)).<br>(ICD-10 PCS) 047Cxxx; 047Dxxx; 047Exxx; 047Fxxx; 047Hxxx; 047Jxxx; 047Kxxx; 047Lxxx; 047Mxxx; 047Nxxx; 047Pxxx; 047Qxxx; 047Rxxx; 047Sxxx; 047Txxx; 047Uxxx; 047Vxxx; 047Wxxx; 047Yxxx (lower extremities, dilation). 04C0xxx; 04CCxxx; 04CDxxx; 04CExxx; 04CFxxx; 04CHxxx; 04CJxxx; 04CKxxx; 04CLxxx; 04CMxxx; 04CNxxx; 04CPxxx; 04CQxxx; 04CRxxx; 04CSxxx; 04CTxxx; 04CUxxx; 04CVxxx; 04CY (extirpation of lower extremity artery). 04100Jx; 041C0Jx; 041D0Jx; 041H0Jx; 041H0Jx; 041H0Kx; 041H4Jx; 041J0Jx; 041J4Jx; 041K0Jx; 041K0Zx; 041L09x; 041L0Kx; 041L0Zx; 041Mxxx; 041Nxxx; 041Sxxx; 041Txxx; 041Uxxx (bypass of lower extremity artery).<br>(CPT) Peripheral Revascularization (carotid and cerebral included) 37205, 37206, 37207, 37208, 37236, 37237, 37184, 37185, 37186, 35302, 35303, 35304, 35305, 35306, 35331, 35351, 35355, 35361, 35363, 35371, 35372, 35381, 35452, 35454, 35456, 35459, 35470, 35472, 35473, 35474, 35483, 35492, 35493, |

| | | |
|---|---|---|
| | | 35495, 35521, 35533, 35537, 35538, 35539, 35540, 35556, 35558, 35563, 35565, 35566, 35571, 35583, 35585, 35587, 35621, 35623, 35637, 35638, 35646, 35647, 35654, 35656, 35661, 35663, 35665, 35666, 35671, 35700, 35876, 35879, 35881, 35883, 35884, 37184, 37185, 37186, 37205, 37206, 37207, 37208, 0236T, 0237T, 0238T, 37225, 37224, 37227, 37226, 37222, 37223, 37220, 37221, 37229, 37228, 37231, 37230, 37233, 37232, 37235, 37234, 35548 |
| | Cerebral vascular disease | (ICD-9) 433.xx (occlusion and stenosis of precerebral arteries); 434.x (occlusion and stenosis of cerebral arteries); 435.x (TIA); 437.x (cerebral atherosclerosis) (ICD-10) G45.x (TIA); I63.xx (cerebral infarction); I65.xx (occlusion and stenosis of precerebral arteries, not resulting in cerebral infarction); I66.xx (occlusion and stenosis of cerebral arteries, not resulting in cerebral infarction); I67.2 (cerebral atherosclerosis) |
| | Myocardial diseases | (ICD-9) 398.xx (other rheumatic heart disease); 402.xx (hypertensive heart disease); 404.xx (hypertensive heart and chronic kidney disease); 415.0 (acute cor pulmonale); 416.8, 416.9 (chronic pulmonary heart disease); 422.xx (acute myocarditis); 425.xx (cardiomyopathy); 428.xx (heart failure); 429.0, 429.1, 429.3, 429.8x (other ill-defined heart diseases). (ICD-10) I11.x (hypertensive heart disease); I13.x (hypertensive heart and chronic kidney disease); I25.3 (aneurysm of heart); I26.0x (pulmonary embolism with acute cor pulmonale); I27.xx (other pulmonary heart disease); I40.x (acute myocarditis); I42.x (cardiomyopathy); I43 (cardiomyopathy in diseases classified elsewhere); I50.xxx (heart failure); I51.xx (complications and ill-defined descriptions of heart disease); I52 (other heart disorders in diseases classified elsewhere). |
| | Pericardial diseases | (ICD-9) 391 (chronic rheumatic pericarditis); 420.xx (acute pericarditis); 423.x (other diseases of the pericardium);<br>(ICD-10) I30.x (acute pericarditis); I31.x (other diseases of the pericardium); I32 (pericarditis in diseases classified elsewhere) |
| | Valvular heart diseases | (ICD-9) 391.x (rheumatic fever with heart involvement); 394.x (diseases of the mitral valve); 395.x (diseases of the aortic valve); 396.x (diseases of the mitral and aortic valve); 397.x (diseases of other endocardial structures); 421.x (acute and subacute |

| | | |
|---|---|---|
| | | endocarditis); 424.xx (other diseases of the endocardium);<br>(ICD-10) I01.x (acute rheumatic fever w/ heart involvement); I05.x (rheumatic mitral valve disease); I06.x (rheumatic aortic valve disease); I07.x (rheumatic tricuspid valve disease); I08.x (multiple valve disease); I09.x (other rheumatic heart diseases); I33.x (acute and subacute endocarditis); I34.x (nonrheumatic mitral valve disorders); I35.x (nonrheumatic aortic valve disorders); I36.x (nonrheumatic tricuspid valve disorders); I37.x (nonrheumatic pulmonary valve disorders); I38 (endocarditis, valve unspecified); I39 (endocarditis and heart valve disorders in diseases classified elsewhere) |
| | Arrhythmias | (ICD-9) 426.xx (conduction disorders); 427.xx (cardiac dysrhythmias) (ICD-10) I44.xx (atrioventricular and left bundle branch block); I45.xx (other conduction disorders); I47.x (paroxysmal tachycardia); I48.xx (atrial fibrillation and flutter); I49.xx (other cardiac arrhythmias.<br>*Procedure*:<br>(ICD-10) 02K8xxx (conduction mapping) |
| | Aortic Disease | (ICD-9) 441.xx (aortic aneurysm and dissection), (ICD-10) I71.xx (aortic<br>aneurysm and dissection) |
| | Congenital heart disease | (ICD-9) 745.xx (bulbus cordis anomalies and anomalies of cardiac septal closure); 746.xx (other congenital anomalies of heart); 747.0x - 747.4x (other congenital anomalies of the circulatory system).<br>(ICD-10) Q20.x (congenital malformations of cardiac chambers and connections); Q21.x (congenital malformations of the cardiac septa); Q22.x (congenital malformations of the pulmonary and tricuspid valves); Q23.x (congenital malformations of the aortic and mitral valves); Q24.x (other congenital malformations of the heart); Q25.xx (congenital malformations of great arteries); Q26.x (congenital abnormalities of great veins) |
| | Other CAD | (ICD-10) I25.4x (coronary artery aneurysm and dissection) |
| Diabetes | (ICD-9) 249.xx (secondary diabetes); 250.xx (diabetes mellitus); 357.2 (diabetic polyneuropathy); 362.0x (diabetic retinopathy); 366.41 (diabetic cataract)<br>(ICD-10) E08.x (diabetes mellitus due to underlying condition), E09.x (drug or chemical- induced diabetes mellitus), E10.x (type 1 diabetes | |

| | | |
|---|---|---|
| | mellitus), E11.x (type 2 diabetes mellitus), E13.x (other specified diabetes mellitus) E14.x (Unspecified diabetes mellitus) | |
| Hypertension | (ICD-9) 401.x (essential hypertension); 403.xx (hypertensive chronic kidney disease); 405.xx (secondary hypertension) (ICD-10) I10 (primary hypertension); I12.x (hypertensive chronic kidney disease); I15.x (secondary hypertension) | |
| Obesity | BMI as provided in MarketScan data (continuous variable) | |
| Chronic Kidney Disease | (ICD-9) 585.3, 585.4 (ICD-10) N18.3x (CKD stage 3); N18.4 (CKD stage 4); N18.5 (CKD stage 5); N18.6 (ESRD) | |
| Hyperlipidemia | (ICD-9) 272 (ICD-10) E78.xx (disorders of lipoprotein metabolism, excluding E78.7 and E78.8); E88.81 (metabolic syndrome) | |
| Tobacco use | Currently using cigars (yes/no), previously used cigars (yes/no), number of cigarettes smoked per day (coded 1-4 based on response), currently using cigarettes (yes/no), number of years of cigarette use (coded 1-4 based on response), packs of cigarettes daily (<1, >1), previous use of cigarettes (yes/no), previous use of chewing tobacco (yes/no), current use of chewing tobacco (yes/no) | |
| Cancer | Lip, oral cavity, pharynx (head and neck) | (ICD-9) 140.0, 140.1, 140.3, 140.4, 140.5, 140.6, 140.8, 140.9, 141.0, 141.1, 141.2, 141.3, 141.4, 141.5, 141.6, 141.8, 141.9, 142.0, 142.1, 142.2, 142.8, 142.9, 143.0, 143.1, 143.8, 143.9, 144.0, 144.1, 144.8, 144.9, 145.0, 145.1, 145.2, 145.3, 145.4, 145.5, 145.6, 145.8, 145.9, 146.0, 146.1, 146.2, 146.3, 146.4, 146.5, 146.6, 146.7, 146.8, 146.9, 147.0, 147.1, 147.2, 147.3, 147.8, 147.9, 148.0, 148.1, 148.2, 148.3, 148.8, 148.9, 149.0, 149.1, 149.8, 149.9 (ICD-10) C00.0, C00.1, C00.2, C00.3, C00.4, C00.5, C00.6, C00.7, C00.8, C00.9, C01, C02.0, C02.1, C02.2, C02.3, C02.4, C02.8, C02.9, C03.0, C03.1, C03.9, C04.0, C04.1, C04.8, C04.9, C05.0, C05.1, C05.2, C05.8, C05.9, C06.0, C06.1, C06.2, C06.8, C06.9, C07, C08.0, C08.1, C08.9, C09.0, C09.1, C09.8, C09.9, C10.0, C10.1, C10.2, C10.3, C10.4, C10.8, C10.9, C11.0, C11.1, C11.2, C11.3, C11.8, C11.9, C12, C13.0, C13.1, C13.2, C13.8, C13.9, C14.0, C14.2, C14.8 |
| | Esophagus | (ICD-9) 150.0, 150.1, 150.2, 150.3, 150.4, 150.5, 150.8, 150.9 (ICD-10) C15.3, C15.4, C15.5, C15.8, C15.9 |
| | Stomach | (ICD-9) 151.0, 151.1, 151.2, 151.3, 151.4, 151.5, 151.6, 151.8, 151.9 (ICD-10) C16.0, C16.1, C16.2, C16.3, C16.4, C16.5, C16.6, C16.8, C16.9 |
| | Small intestine | (ICD-9) 152.0, 152.1, 152.2, 152.3, 152.8, 152.9 (ICD-10) C17.0, C17.1, C17.2, C17.3, C17.8, C17.9, C26.0 |

| | | |
|---|---|---|
| | Colon | (ICD-9) 153.0, 153.1, 153.2, 153.3, 153.4, 153.5, 153.6, 153.7, 153.8, 153.9 (ICD-10) C18.0, C18.1, C18.2, C18.3, C18.4, C18.5, C18.6, C18.7, C18.8, C18.9 |
| | Rectosigmoid, rectum, anus | (ICD-9) 154.0, 154.1, 154.2, 154.3, 154.8 (ICD-10) C19, C20, C21.0, C21.1, C21.2, C21.8 |
| | Liver, gallbladder, spleen | (ICD-9) 155.0, 155.1, 155.2, 156.0, 156.1, 156.2, 156.8, 156.9, 159.0, 159.1 (ICD-10) C22.0, C22.1, C22.2, C22.3, C22.4, C22.7, C22.8, C22.9, C23, C24.0, C24.1, C24.8, C24.9, C26.1, C26.9 |
| | Pancreas | (ICD-9) 157.0, 157.1, 157.2, 157.3, 157.4, 157.8, 157.9 (ICD-10) C25.0, C25.1, C25.2, C25.3, C25.4, C25.7, C25.8, C25.9 |
| | Nasal cavity, middle ear, accessory sinuses | (ICD-9) 160.0, 160.1, 160.2, 160.3, 160.4, 160.5, 160.8, 160.9 (ICD-10) C30.0, C30.1, C31.0, C31.1, C31.2, C31.3, C31.8, C31.9 |
| | Larynx, trachea | (ICD-9) 161.0, 161.1, 161.2, 161.3, 161.8, 161.9, 162.0 (ICD-10) C32.0, C32.1, C32.2, C32.3, C32.8, C32.9, C33 |
| | Lung | (ICD-9) 162.2, 162.3, 162.4, 162.5, 162.8, 162.9, 165.0, 165.8, 165.9 (ICD-10) C34.0, C34.1, C34.2, C34.3, C34.8, C34.9, C39.0, C39.9 |
| | Thymus, heart, mediastinum, pleura | (ICD-9) 163.0, 163.1, 163.8, 163.9, 164.0, 164.1, 164.2, 164.3, 164.8, 164.9 (ICD-10) C37, C38.0, C38.1, C38.2, C38.3, C38.4, C38.8 |
| | Bone and articular cartilage | (ICD-9) 170.0, 170.1, 170.2, 170.3, 170.4, 170.5, 170.6, 170.7, 170.8, 170.9 (ICD-10) C40.0, C40.1, C40.2, C40.3, C40.8, C40.9, C41.0, C41.1, C41.2, C41.3, C41.4, C41.9 |
| | Melanoma | (ICD-9) 172.0, 172.1, 172.2, 172.3, 172.4, 172.5, 172.6, 172.7, 172.8, 172.9 (ICD-10) C43.0, C43.1, C43.2, C43.3, C43.4, C43.5, C43.6, C43.7, C43.8, C43.9 |
| | Mesothelial and soft tissue | (ICD-9) 158.0, 158.8, 158.9, 171.0, 171.2, 171.3, 171.4, 171.5, 171.6, 171.7, 171.8, 171.9, 176.0, 176.1, 176.2, 176.3, 176.4, 176.5, 176.8, 176.9 (ICD-10) C45.0, C45.1, C45.2, C45.7, C45.9, C46.0, C46.1, C46.2, C46.3, C46.4, C46.5, C46.7, C46.9, C47.0, C47.1, C47.2, C47.3, C47.4, C47.5, C47.6, C47.8, C47.9, C48.0, C48.1, C48.2, C48.8, C49.0, C49.1, C49.2, C49.3, C49.4, C49.5, C49.6, C49.8, C49.9, C49.A |
| | Breast | (ICD-9) 174.0, 174.2, 174.3, 174.4, 174.5, 174.6, 174.8, 174.9, 175.0, 175.9 (ICD-10) C50.0, C50.1, C50.2, C50.3, C50.4, C50.5, C50.6, C50.8, C50.9 |
| | Uterine | (ICD-9) 179, 182.0, 182.1, 182.8 |

| | | |
|---|---|---|
| | | (ICD-10) C54.0, C54.1, C54.2, C54.3, C54.8, C54.9, C55 |
| | Ovarian | (ICD-9) 183 (ICD-10) C56.1, C56.2, C56.9 |
| | Female genital organs, other | (ICD-9) 180.0, 180.1, 180.8, 180.9, 181, 183.2, 183.3, 183.4, 183.5, 183.8, 183.9, 184.0, 184.1, 184.2, 184.3, 184.4, 184.8, 184.9<br>(ICD-10) C51.0, C51.1, C51.2, C51.8, C51.9, C52, C53.0, C53.1, C53.8, C53.9, C57.0, C57.1, C57.2, C57.3, C57.4, C57.7, C57.8, C57.9, C58 |
| | Prostate | (ICD-9) 185 (ICD-10) C61, Z19.1, Z19.2 |
| | Male genital organs, other | (ICD-9) 186.0, 186.9, 187.1, 187.2, 187.3, 187.4, 187.5, 187.6, 187.7, 187.8, 187.9 (ICD-10) C60.0, C60.1, C60.2, C60.8, C60.9, C62.0, C62.1, C62.9, C63.0, C63.1, C63.2, C63.7, C63.8, C63.9 |
| | Renal cell carcinoma | (ICD-9) 189.0, 189.1<br>(ICD-10) C64.1, C64.2, C64.9, C65.1, C65.2, C65.9, C66.1, C66.2, C66.9 |
| | Bladder | (ICD-9) 188.0, 188.1, 188.2, 188.3, 188.4, 188.5, 188.6, 188.7, 188.8, 188.9 (ICD-10) C67.0 , C67.1, C67.2, C67.3, C67.4, C67.5, C67.6, C67.7, C67.8, C67.9 |
| | Urinary organs, other | (ICD-9) 189.2, 189.3, 189.4, 189.8, 189.9<br>(ICD-10) C68.0, C68.1, C68.8, C68.9 |
| | Eye and adnexa | (ICD-9) 190.0, 190.1, 190.2, 190.3, 190.4, 190.5, 190.6, 190.7, 190.8, 190.9 (ICD-10) C69.0, C69.1, C69.2, C69.3, C69.4, C69.5, C69.6, C69.8, C69.9 |
| | Brain, meninges, nerves | (ICD-9) 191.0, 191.1, 191.2, 191.3, 191.4, 191.5, 191.6, 191.7, 191.8, 191.9, 192.0, 192.1, 192.2, 192.3, 192.8, 192.9<br>(ICD-10) C70.0, C70.1, C70.9, C71.0, C71.1, C71.2, C71.3, C71.4, C71.5, C71.6, C71.7, C71.8, C71.9, C72.0, C72.1, C72.2, C72.3, C72.4, C72.5, C72.9 |
| | Thyroid, endocrine (other) | (ICD-9) 193, 194.0, 194.1, 194.3, 194.4, 194.5, 194.6, 194.8, 194.9 (ICD-10) C73, C74.0, C74.1, C74.9, C75.0, C75.1, C75.2, C75.3, C75.4, C75.5, C75.8, C75.9 |
| | Neuroendocrine tumors | (ICD-9) 209.0, 209.1, 209.2, 209.3, 209.4, 209.5, 209.6, 209.7 (ICD-10) C7A.0, C7A.1, C7A.8, C7B.0, C7B.1, C7B.8 |
| | Lymphoma (B and T), plasma cell dyscrasias | (ICD-9) 200.0, 200.1, 200.2, 200.3, 200.4, 200.5, 200.6, 200.7, 200.8, 201.0, 201.1, 201.2, 201.4, 201.5, 201.6, 201.7, 201.9, 202.0, 202.1, 202.2, 202.3, 202.4, 202.5, 202.7, 202.8, 202.9, 203.0, 203.1<br>(ICD-10) C81.0, C81.1, C81.2, C81.3, C81.4, C81.7, C81.9, C82.0, C82.1, C82.2, C82.3, C82.4, C82.5, C82.6, C82.8, C82.9, C83.0, C83.1, C83.3, C83.5, |

| | | |
|---|---|---|
| | | C83.7, C83.8, C83.9, C84.0, C84.1, C84.4, C84.6, C84.7, C84.A, C84.Z, C84.9, C85.1, C85.2, C85.8, C85.9, C86.0, C86.1, C86.2, C86.3, C86.4, C86.5, C86.6, C88.0, C88.2, C88.3, C88.4, C88.8, C88.9, C90.0, C90.1, C90.2, C90.3 |
| | Leukemia (lymphoid, myeloid, and other) | (ICD-9) 204.0, 204.1, 204.2, 204.8, 204.9, 205.0, 205.1, 205.2, 205.3, 205.8, 205.9, 206.0, 206.1, 206.2, 206.8, 206.9, 207.0, 207.1, 207.2, 207.8, 208.0, 208.1, 208.2, 208.8, 208.9<br>(ICD-10) C91.0, C91.1, C91.3, C91.4, C91.5, C91.6, C91.A, C91.Z, C91.9, C92.0, C92.1, C92.2, C92.3, C92.4, C92.5, C92.6, C92.A, C92.Z, C92.9, C93.0, C93.1, C93.3, C93.Z, C93.9, C94.0, C94.2, C94.3, C94.4, C94.6, C94.8, C95.0, C95.1, C95.9 |
| | Hematologic malignancies, other | (ICD-9) 202.6, 203.8, 238.4, 238.7<br>(ICD-10) C96.0, C96.2, C96.4, C96.5, C96.6, C96.A, C96.Z, C96.9, D45, D46 |
| | Personal history of malignancy (only used for exclusion but not for incident cancer) | (ICD-9) V10.x (excluding V10.83), V87.41, V87.43<br>(ICD-10) Z85.x (excluding Z85.82), Z86.00x, Z92.21, Z92.23 |

**Table S8. ICD codes for UK Biobank**

| Category | Subcategory | UKB Field | UKB/ICD Code (Include) | UKB/ICD Code (Exclude) | Notes |
|---|---|---|---|---|---|
| Cancer | Any cancer | 40006 | Chapter II (Neoplasms) | | n=152,636 |
| | Colorectal cancer | 40006 | C18 (Malignant neoplasm of colon) | | n=6,245 |
| CVD | Myocardial infarction | 41270 | I21, I22 | | n=19,845; 1,026 |
| | Coronary artery disease | 41270 | I21, I22, I23, I24, I25 | I253, I254 | Exclude heart/ coronary aneurysm |
| | Peripheral artery disease / Aortic atherosclerosis | 41270 | I65, I70, I739, I75 | | I75 empty; I65 overlap with cerebrovascular |
| | Cerebrovascular disease | 41270 | G45, I63, I65, I66, I672 | | I65 overlap with PAD |
| | Myocardial diseases | 41270 | I11, I13, I253, I26.0, I27, I40, I42, I43, I50, I51, I52 | | |
| | Pericardial diseases | 41270 | I30, I31, I32 | | |
| | Valvular heart diseases | 41270 | I01, I05, I06, I07, I08, I09, I33, I34, I35, I36, I37, I38, I39 | | |
| | Aortic disease | 41270 | I71 | | |
| | Congenital heart disease | 41270 | Q20, Q21, Q22, Q23, Q24, Q25, Q26 | | |
| | Other CAD | 41270 | I25.4 | | Coronary artery aneurysm |
| Risk factors | Diabetes | 41270 | E08, E09, E10, E11, E13, E14 | | E08/E09 may be empty |
| | Hypertension | 41270 | I10, I12, I15 | | n=162,262; 2,408; 286 |
| | Chronic kidney disease | 41270 | N183, N184, N185, N186 | | N186 (ESRD) may be empty |

| | | | | | |
|---|---|---|---|---|---|
| | Hyperlipidemia | 41270 | E78, E88.8 | E787, E788 | Exclude bile acid / other lipid disorders |
| | Smoking status | 22506 | 111 (most/all days), 112 (occasionally), 113 (ex-smoker) | | |

**Table S9. Demographics of patient cohorts**

| Characteristic | Colorectal cancer | Prostate cancer |
|---|---|---|
| Overall | 6908 | 11929 |
| Age Category (at time of surgery) | | |
| 18-39 | 345 | 17 |
| 40-49 | 614 | 476 |
| 50-59 | 1349 | 3565 |
| 60-64 | 732 | 2990 |
| 65-69 | 615 | 2159 |
| 70-79 | 1459 | 1394 |
| 80+ | 1254 | 134 |
| Sex | | |
| Male | 3369 | 11929 |
| Female | 3539 | 0 |

**Table S10. Diagnostic and procedural codes**

| | | |
|---|---|---|
| **Colorectal cancer** | Colon | ICD9:153, 153.0, 153.1, 153.2, 153.3, 153.4, 153.5, 153.6, 153.7, 153.8, 153.9; ICD10:C18, C18.0, C18.1, C18.2, C18.3, C18.4, C18.5, C18.6, C18.7, C18.8, C18.9 |
| | Rectosigmoid junction | ICD9:154.0; ICD10:C19 |
| | Rectum | ICD9:154.1; ICD10:C20 |
| **Prostate cancer** | | ICD9:185; ICD10:C61 |
| **PCI** | CPT | 92920, 92921, 92924, 92925, 92928, 92929, 92933, 92934, 92937, 92938, 92941, 92943, 92944, C9600, C9601, C9602, C9603, C9604, C9605, C9606, C9607, C9608 |
| | EPIC | 27000145, 27000146, 27000147, 48000103, 48000234, 48000235, 48000236, 48000238, 48000239, 48000240, 48000242, 48000244, 48000245, 48000246, 48000327, 48000404, 48000405, 48000406, 48000407, 48000408, 48000409, 48000410, 48000411, 62400161 |
| | ICD9 | 00.66, 17.55, 36.01, 36.02, 36.05, 36.06, 36.07 |
| | ICD10 | 0270346, 027034Z, 027035Z, 027036Z, 027037Z, 02703D6, 02703DZ, 02703TZ, 02703Z6, 02703ZZ, 027044Z, 02704DZ, 0271346, 027134Z, 0271356, 027135Z, 027136Z, 027137Z, 02713DZ, 02713ZZ, 027234Z, 027236Z, 027237Z, 027334Z |
| | HCPCS | C9600, C9604, C9606, G0290, G0291 |
| | ONCALL | MLFL4, MLFL4-3, MXAB2, MYRD1 |
| **CABG** | CPT | 33510, 33511, 33512, 33513, 33514, 33516, 33517, 33518, 33519, 33520, 33521, 33522, 33523, 33525, 33528, 33530, 33533, 33534, 33535, 33536, 35600 |
| | ICD9 | 36.11, 36.12, 36.13, 36.14, 36.15, 36.16, 36.19, 36.2, 36.31 |
| | ICD10 | 0210093, 0210098, 0210099, 021009W, 02100A3, 02100A8, 02100A9, 02100AC, 02100AW, 02100KW, 02100Z8, 02100Z9, 02100ZC, 0211093, 021109W, 02110AW, 02110Z3, 02110Z9, 0212093, 021209W, 02120ZC, 0213093, 021309W |
| | LMR | LPA497 |
| | ONCALL | BLER5, BLER5-3, MYRD1, MYRD1-1 |
